# Blocking neuroestrogen synthesis modifies neural representations of learned song without altering vocal imitation accuracy in developing songbirds

**DOI:** 10.1101/702704

**Authors:** Daniel M. Vahaba, Amelia Hecsh, Luke Remage-Healey

## Abstract

Birdsong, like human speech, is learned early in life by first memorizing an auditory model. Once memorized, birds compare their own burgeoning vocalizations to their auditory memory, and adjust their song to match the model. While much is known about this latter part of vocal learning, less is known about how initial auditory experiences are formed and consolidated. In both adults and developing songbirds, there is strong evidence suggesting the caudomedial nidopallium (NCM), a higher order auditory forebrain area, is the site of auditory memory consolidation. However, the mechanisms that facilitate this consolidation are poorly understood. One likely mechanism is 17β-estradiol (E2), which is associated with speech-language development and disorders in humans, and is abundant in both mammalian temporal cortex and songbird NCM. Circulating E2 is also elevated during the auditory memory phase, and in NCM immediately after song learning sessions, suggesting it functions to encode recent auditory experience. Therefore, we tested a role for E2 production in auditory memory consolidation during development using a comprehensive set of investigations to ask this question at the level of neuroanatomy, neurophysiology, and behavior. Our results demonstrate that while systemic estrogen synthesis blockade regulates juvenile song production, inhibiting E2 synthesis locally within NCM does not adversely affect song learning outcomes. Surprisingly, early life E2 manipulations in NCM modify the neural representations of birds’ own song and the model tutor song in both NCM and a downstream sensorimotor nucleus (HVC). Further, we show that the capacity to synthesize neuroestrogens remains high throughout development alongside substantial changes in NCM cell density across age. Taken together, these findings suggest that E2 plays a multifaceted role during development, and demonstrate that contrary to prediction, unilateral post-training estrogen synthesis blockade in the auditory cortex does not negatively impact vocal learning. Acute downregulation of neuroestrogens are therefore likely permissive for juvenile auditory memorization, while neuroestrogen synthesis influences communication production and representation in adulthood.

## INTRODUCTION

While many animals use sounds to communicate with one another (vocal communication), the ability to learn to vocally communicate is relatively rare (Petkov and Jarvis, 2012). In vocal learning animals, such as humans and songbirds, vocal learning occurs across two main phases: an auditory memorization (‘sensory’) phase, followed by a sensorimotor phase (‘babbling’, error correction/feedback) (Kuhl, 2010; Derégnaucourt, 2011). While much is known about sensorimotor learning, the mechanisms that guide the formation and consolidation of auditory memories early in life are less clear.

One brain region likely involved in storing auditory memories in songbirds is the caudomedial nidopallium (NCM) (Bolhuis and Moorman, 2015). NCM, comparable to mammalian secondary auditory cortex, is required for accurate song learning. Blocking ERK-signaling bilaterally in NCM during tutoring leads to poor song imitation (London and Clayton, 2008). Tutoring naïve juvenile songbirds leads to an increased proportion of tutor-song-selective neurons in NCM (Yanagihara and Yazaki-Sugiyama, 2016). Further, bilateral NCM lesions abolish innate preference for tutor song in adults (Gobes and Bolhuis, 2007; but see Canopoli et al., 2016; Canopoli et al., 2017). Thus, NCM contains a putative tutor ‘engram’; however, the neuromodulatory mechanisms that shape this auditory memory formation and consolidation remain unknown.

Consolidating recent experience in other contexts and systems require presynaptic signaling molecules (‘neuromodulators’), such as brain-derived estrogens. The predominant estrogen 17β-estradiol (E2) is a candidate neuromodulator required for auditory memory consolidation, as is evident in a similar role in adult hippocampal-dependent cognition, across taxa (Vierk et al., 2012; Srivastava et al., 2013; Luine, 2014) (Woolley and McEwen, 1992; Packard and Teather, 1997; Packard, 1998; Woolley, 2007; Frick, 2012; Bailey et al., 2013; Rensel et al., 2013; Bailey and Saldanha, 2015; Rensel et al., 2015; Barth et al., 2016; Tuscher et al., 2016; Bailey et al., 2017; Blaustein, 2017; Bayer et al., 2018), but see (Korol and Pisani, 2015). Additionally, both circulating and brain-derived estrogens (‘neuroestrogens’) typically enhance hearing (Caras, 2013; Caras and Remage-Healey, 2016), and are associated with language and verbal memory in humans (Fernandez et al., 2003; Zimmerman et al., 2011; Anthoni et al., 2012; Wermke et al., 2014; Schaadt et al., 2015). Together, current evidence suggests that neuroestrogen signaling may facilitate the consolidation of recent auditory experience (Vahaba and Remage-Healey, 2018).

Neuroestrogens rapidly enhance the gain and coding of auditory neurons in NCM across the lifespan (Remage-Healey et al., 2010; Remage-Healey and Joshi, 2012; Vahaba et al., 2017). Thus, it is possible that one functional role of E2 acting within the auditory forebrain is to facilitate song memory consolidation. NCM is uniquely enriched with estrogen synthase (aromatase) in vocal learning birds (Silverin et al., 2000), suggesting its presence is distinctly important for song learning. Further, systemic inhibition of estrogen synthesis during training and testing results in impaired auditory recognition in adult zebra finches (Yoder et al., 2012). However, it is unclear how neuroestrogens may affect song learning during tutor memorization. Currently, there are several pieces of evidence that suggest a functional role for E2 during the vocal learning critical period. In songbirds, circulating E2 levels rise during the sensory phase, and at least in swamp sparrows, predict future tutor imitation success (Pröve, 1983; Weichel et al., 1986; Marler et al., 1987; Marler et al., 1988), as in humans with language (Wermke et al., 2014; Quast et al., 2016). Moreover, the expression of GPER1 (a membrane-bound estrogen receptor proposed to mediate the rapid effects of E2) peaks in the telencephalon of male songbirds during the sensory phase (Acharya and Veney, 2011). As with E2-dependent learning in rodents, E2 levels are rapidly elevated in NCM immediately after a song learning session (Chao et al., 2015).

The aim of the present study was to determine whether E2 synthesis supports the consolidation of a recent auditory experience and the eventual vocal imitation of a tutor model. Based on prior findings, we postulated that elevated E2 levels in the auditory forebrain aid in memory consolidation following individual song learning bouts. We tested whether the eventual degree of vocal similarity between the social model (tutor) and the pupil in adulthood would be impaired by inhibiting neuroestrogen synthesis in NCM during and immediately after bouts of vocal communication learning.

## MATERIALS & METHODS

All methods and procedures were approved by the Institutional Animal Care and Use Committee (IACUC) at the University of Massachusetts Amherst.

### Immunocytochemistry

#### Animals, perfusion, and sectioning

We first sought to confirm the presence of aromatase in NCM across development. While previous studies have assessed aromatase expression in the brains of developing songbird (Vockel et al., 1988; Jacobs et al., 1999; Saldanha et al., 1999; Saldanha et al., 2000; Chao et al., 2015), there is limited information on aromatase protein expression within NCM between the sensory and sensorimotor periods, which represents a critical transition in auditory response and neuroestrogen sensitivity (Vahaba et al., 2017). Male juvenile zebra finches (*n* = 6) were selected from mixed-sex breeding aviaries maintained on a 14:10 light:dark cycle. Male sensorimotor subjects (*n =* 3; 65, 71, and 71 dph) were identified by their sexually dimorphic plumage (orange cheek feathers; brown and black badge feathers). Sensory-aged male subjects without dimorphic plumage (*n* = 3; 20, 26, and 34 dph) were identified by PCR (see *Sex Determination* below). All subjects were obtained from our breeding colony and were exposed to adult song up until the day of the perfusion. Birds were euthanized via anesthetic overdose (isoflurane) and transcardially perfused with 20 - 30 mL of 0.1M phosphate buffer saline (PBS) followed by 35 mL of 4% paraformaldehyde (PFA). After perfusion, brains were extracted and placed into 4% PFA for 24 hours at 4° C. Brains were then transferred to a 30% sucrose-0.1 M PBS solution for 24 – 48 hours at 4° C. Once fixed, brains were submerged in an opaque tissue-embedding medium (O.C.T. compound; Tissue-Plus; Fisher Health-Care) and frozen at -80° C. Brains were thawed on wet ice on the day of sectioning and hemisected using a razor blade to allow us to carefully distinguish hemispheres. Brains were sectioned at 35 µm in the sagittal plane at -20° C using a cryostat (Leica CM3050 S). Each hemisphere was separately collected into two series for lateral sections, and four series for medial sections. Medial sections were determined by the emergence of cerebellum. Sectioned tissue was placed in cryoprotectant medium in 12-well plates, which was wrapped with Parafilm and stored at -20° C until immunocytochemistry.

#### Antibodies

Antibodies and dilutions for aromatase and parvalbumin were identical to those used in Ikeda et al. (2017). Briefly, we used a polyclonal anti-aromatase primary antibody raised in rabbit (1:2,000; a generous gift from Dr. Colin Saldanha), and a monoclonal anti-parvalbumin primary antibody raised in mouse (1:10,000; Millipore MAB1572; RRID: AB_2174013). Secondary antibodies included goat anti-rabbit Alexa 488 (1:500; Thermo Fisher Scientific Inc.), and goat anti-mouse Alexa 647 (1:100; Thermo Fisher Scientific Inc.).

#### Procedure

Brain sections were first manually washed 3 x in 0.1 M PB, followed by 3 x 15-minute washes in 0.1 M PB on a plate shaker, followed by a 2-hour incubation at room temperature with 10% normal goat serum (Vector) in 0.3 % PBT. Tissue was then transferred to a 10% normal goat serum-0.3% PBT solution containing the primary antibodies and incubated at room temperature for 60 minutes. Afterwards, plates were tightly wrapped in parafilm and placed on an orbital shaker in a cold room at 4° C for 48 hours. On day 3, tissue was washed 3 x 15 minutes in 0.1 % PBT before being transferred to the secondary antibody-containing solution made in 0.3% PBT for 60 minutes. At this point, tissue was kept in the dark to prevent any florescent bleaching. Tissue was washed again 3 x 10 minutes in 0.1% PBT, and finally transferred to 0.1 M PB, wrapped in parafilm, and stored at 4° C. Several days later, tissue was slide mounted, covered with ProLong Diamond Antifade Mountant with DAPI (Thermo Fisher Scientific Inc.), cover slipped, and placed in an opaque slide box and stored at 4° C.

#### Confocal imaging

Fluorescently-labelled tissue was imaged using a confocal microscope (Nikon A1 Resonant Confocal) with NIS-Elements imaging software. The laser strength and gain were determined independently for each antibody/fluorescent channel of interest. Once the levels were determined, the same setting was applied across all sections per fluorescent channel. NCM was located anatomically by the presence of cerebellum and the absence of aromatase-rich nucleus taenia (TnA; lateral boundary of NCM). An overview/reference image at 10x was obtained for each section followed by subregion (dNCM and vNCM) z-stacks obtained at 60x (1 µm z-steps for 15 µm).

#### Image analysis

An experimenter blind to subjects’ ages and hemisphere quantified the total number immunostained cells for each fluorescent channel using ImageJ 1.52h (Schneider et al., 2012). We measured immunopositive-neurons two ways. Initially, we quantified aromatase and parvalbumin immunopositive-cells by calculating their expression as a percentage of DAPI to normalize for relative cell density across sections and subjects (e.g. Aromatase+ cells % of DAPI = total # of aromatase+ cells / total # of DAPI+ cells). Additionally, we also quantified cell density relative to image volume to provide a more standardized report of its expression using the following equation:

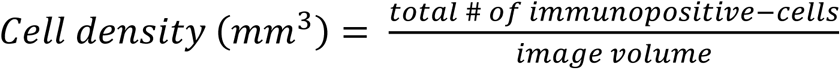

### Juvenile Song Learning

#### Animals

Juvenile male zebra finches (*Taeniopygia guttata*) were obtained from our breeding aviaries (*N* = 34; *n*= 6 for systemic experiments; *n* = 28 for microdialysis experiments. See ***Fig. 1*** for the experimental timeline). Nest boxes with an active clutch of young zebra finches (<10 dph) were observed to identify the putative mother. Once identified, the mother, offspring, and their nest box were placed in a cage within a sound-attenuation chamber (Eckel Acoustics), either as a single-family group, or, in a few rare instances, two adjacent cages of females with siblings were placed in the same chamber. Some breeding pairs were also isolated before laying a clutch (*n* = 3). In these instances, the adult male was left in the sound-attenuation chamber until the fledglings were ∼13 dph. The remaining fledgling were removed from the breeding colony by 13 dph (range = 5 – 17 dph), which is well before the putative opening of the critical period for song learning (∼20 – 25 dph). Birds were confirmed to be male via sex determination PCR at ∼22 dph. By ∼30 dph, most birds were isolated from their siblings and mom (range = 29 – 39 dph; *n* = 2 birds that were >38 dph; most birds were 29 – 31 dph) and placed in a new cage and sound-attenuation chamber along with an unrelated adult companion female. An omnidirectional microphone (Countryman) was placed in the chamber and song was continuously recorded for the remainder of the experiment using Sound Analysis Pro (Tchernichovski et al., 2000).

**Fig. 1.**
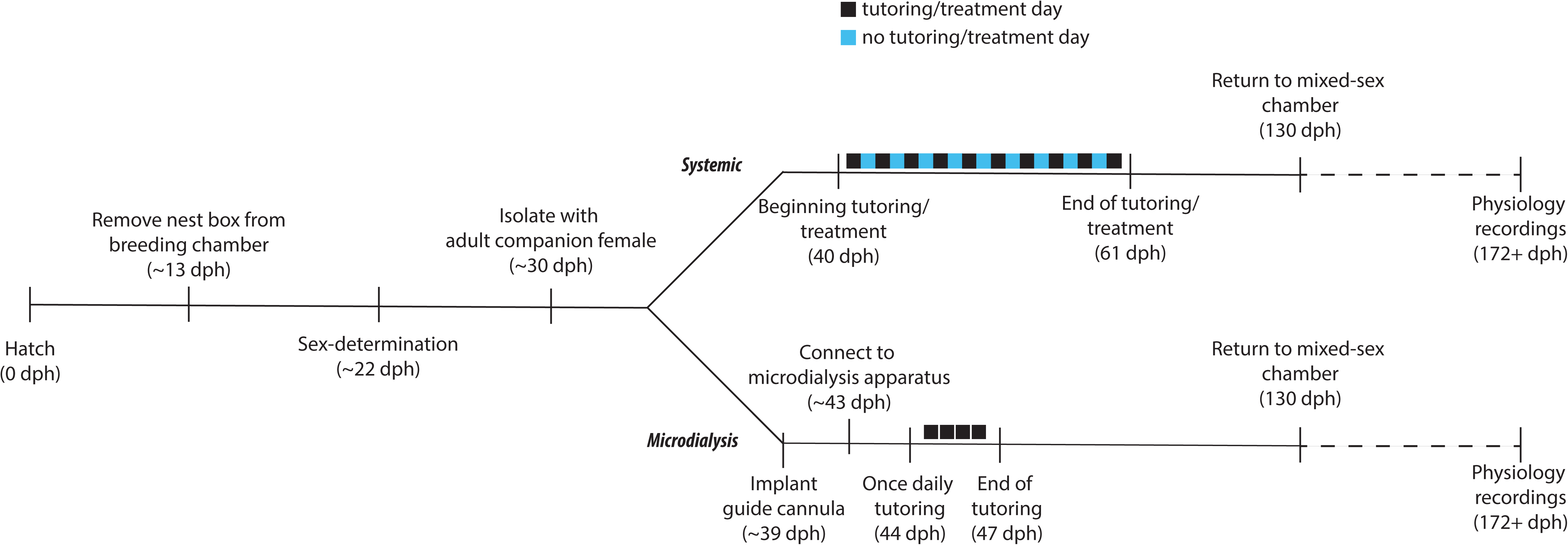
Experimental timeline.

For microdialysis subjects, a total of 20 birds were successfully treated with FAD or aCSF (*n* = 5 subjects per hemisphere per treatment). An additional eight subjects experienced non-health related technical issues during microdialysis (e.g. clogged microdialysis probe) that resulted in them being prematurely disconnected but retained as surgery control subjects (‘cannula’-only subjects). One of these failed microdialysis subjects was deprived of any tutoring or adult male song until after 131 dph and served as an isolate control subject.

### Timeline

#### Systemic

For systemically-treated subjects, birds were co-housed with an adult companion female throughout the entire experiment. Tutoring began at 40 dph (see *Tutoring regiment*) and was immediately followed by oral administration of the assigned treatment. Tutoring continued every other day for 20 days (i.e. 10 days of total tutoring), ending at 60 dph. Peripheral FAD treatment suppresses E2 synthesis for up to 48 hours (Wade et al., 1994). Thus, there was one ‘washout’ day without any treatments between each tutoring session. Birds were returned to group housing at 131 dph, and after at least 6 weeks (∼196 dph), were re-captured to record song and terminal electrophysiology recordings.

#### Microdialysis

Guide cannulae were unilaterally implanted in NCM several days after being initially isolated with a companion female. Several days following surgery, birds were connected to the microdialysis apparatus in a new sound-attenuation chamber without any companion birds. One day later, daily tutoring began for two to three days. After the last tutor session, birds were disconnected from the microdialysis setup and placed in a sound-attenuation chamber with an adult female companion bird in an adjacent cage. Companion females were switched every two weeks. Birds were returned to a group setting (all-male aviary in breeding room, or in a mixed-sex sound-attenuation chamber in same-sex cage) at 131 dph. After at least 6 weeks had elapsed, birds were returned to a sound-attenuation chamber for follow-up song recording and subsequent electrophysiology experiments. After electrophysiology recordings, birds were sacrificed, and brains were extracted for future sectioning and histological examination.

#### Sex determination

Zebra finches begin to develop sexually dimorphic plumage at ∼30 - 40 dph. Therefore, we used established methods (Griffiths et al., 1998) as we have previously described (Chao et al., 2015; Vahaba et al., 2017) to determine juvenile birds’ sex. Briefly, DNA for sex determination PCR was extracted from whole blood obtained from the ulnar vein typically at ∼22 dph (median age = 22 dph; range = 18 – 30 dph). Identified males were retained for the experiment, whereas females were returned to their original breeding aviary along with their mother once the youngest male fledgling reached ∼30 dph.

#### Pharmacological inhibition of aromatase

For systemic experiments, birds were fed 30 µL of either saline (0.9% NaCl in _dd_H20) or FAD (1 mg/mL in 0.9% NaCl) immediately following tutoring cessation. This dose is similar to previous studies that demonstrate significantly reduced aromatase activity and/or estradiol levels in zebra finches (Wade et al., 1994; Saldanha et al., 2000; Saldanha et al., 2004; Remage-Healey et al., 2010; Rensel et al., 2013). Microdialysis subjects were retrodialyzed with artificial cerebrospinal fluid (aCSF) and 100 µM FAD in aCSF prepared as in previous experiments (Remage-Healey et al., 2008; Remage-Healey et al., 2010; Remage-Healey et al., 2012; Chao et al., 2015).

### *In Vivo* Microdialysis

#### Surgery

A unilateral CMA-7 microdialysis guide cannula with obdurator (CMA Microdialysis, CMA 7, ref. no. P000138) was implanted several days after isolation with a companion female (median age = 39 dph; range = 35 – 47 dph), as in previous studies (Remage-Healey et al., 2008; Ikeda et al., 2014; Chao et al., 2015). Birds were food deprived 30 minutes prior to surgery, and then received an intramuscular injection of Equithesin (30 – 40 µL, typically). Twenty minutes later, birds were swaddled in a Kim wipe, and placed atop a heating pad and secured via ear bars at 45° to our custom surgical stereotaxic apparatus (Herb Adams Engineering). Head feathers were removed and a 20 µL subcutaneous injection of 2% lidocaine was administered underneath the scalp, which was subsequently resected to expose the outer layer of skull. The midsagittal sinus bifurcation (MSB) was then identified and used as a 0-point anatomical reference. A unilateral fenestra was then made over one hemisphere of NCM (coordinates: rostral = 1.20 mm, lateral = ± 0.90 mm), and the dura was carefully resected. A CMA-7 guide cannula with obdurator was then descended approximately 1.0 mm ventral into the proximate region of NCM (ventral range of NCM at this coordinate is 0.80 – 1.40 mm). The guide cannula was secured using cyanoacrylate and dental cement, and the exposed scalp and incision area sealed with cyanoacrylate. Birds recovered on a heating pad in a cage with *ad libitum* food and water until awake, after which they were transferred back to their sound-attenuation chamber in a separate cage from the companion female.

Acute neural injury induces glial aromatase production in birds, with aromatase responses peaking at 72 hours, and persisting up to six weeks after insult (Peterson et al., 2004; Wynne et al., 2008; Balthazart and Ball, 2013). To reduce the confound of injury-induced aromatase upregulation from the guide cannula surgery, birds were given at least three days to recover prior to starting microdialysis (median = 4 days; range = 3 – 5 days) to allow for injury-induced glial aromatase levels to subside (Saldanha et al., 2013).

#### Microdialysis

After the recovery period, birds were connected to the microdialysis apparatus in a new sound-attenuation chamber. The obdurator was replaced with a CMA-7 microdialysis probe (1 mm membrane length, CMA Microdialysis, ref. no. P000082), which was then connected to a dual-channel microdialysis swivel (375/D/22QM; Instech Labs) via fluorinated ethylene propylene (FEP) inlet and outlet tubing. Once the bird was connected, aCSF was dialyzed at a rate of 2 µL/min by a syringe pump located outside of the chamber (PHD 1000, Harvard Apparatus). After being hooked-up, all birds were observed to ensure they were healthy as evidenced by eating, drinking, and the ability to comfortably navigate the cage. Dialysate samples were collected every hour during the day (∼09:00 -∼18:00 pm), yielding ∼120 µL of dialysate per sample. Perfusate was dialyzed at a rate of 2 µL/min for the entire duration of the microdialysis experiment. Several hours after the final tutor session, FEP tubing was disconnected and birds were returned to a sound-attenuation chamber in a separate cage alongside an adult companion female. As described in similar studies (London and Clayton, 2008), guide cannulae eventually detach after experiments as the skull develops and expands, without any obvious deleterious health effects, typically 12 days after the last day of microdialysis (range = 6 – 38 days post-final microdialysis day; in one case, this did not occur until 154 days after microdialysis).

#### Tutoring regimen

All birds were naïve to song before the tutoring period. After tutoring, all birds were returned to an individual sound-attenuation chamber with an unrelated adult female companion in an adjacent cage. Including a companion female is atypical for most experimental studies of song learning in the lab, and there is some evidence to suggest that adult females may impact song development in juvenile male zebra finches (Kojima and Doupe, 2011) and cowbirds (King et al., 2005). However, we opted to include a companion female as isolating subjects is less naturalistic for zebra finches (a highly gregarious songbird), and likely a great deal more stressful for developing subjects.

#### Passive audiovisual tutoring playback

In an initial pilot experiment, we were curious whether an automated passive playback tutoring design would enable accurate song learning/imitation in adulthood, as used in other song tutoring studies (Deshpande et al., 2014; Chao et al., 2015). Similar early isolation procedures as with the systemic and microdialysis subjects were used on a separate set of birds (*n* = 8). Otherwise unmanipulated subjects were isolated from their mother and siblings ∼37 dph, and daily tutoring began at 42 dph until 47 dph (5 sessions total). Tutoring began at ∼10:00 each day and lasted for one hour. During the tutoring session, a 60-minute tutoring video was played on a USB-powered LCD monitor (Lilliput 7-in) alongside song broadcasted via an adjacent speaker (Sony; model # SRS-TP1WHI). The video and song were obtained from an adult male zebra finch singing directed song to a female. At 48 dph, birds were reunited with an adult female companion and kept in isolation until 111 dph, after which time they were returned to a mixed-sex aviary. Song was recorded throughout the entirety of the experiment. Overall, birds tutored with passive audiovisual methods produced poor copies of the tutor song (*n* = 6; mean ± SEM; similarity = 41.09% ± 0.07%; range = 23.37% - 64.98%), likely due to zebra finches requiring active/self-solicited learning (e.g. operant tutoring) and/or social instruction (reviewed in Derégnaucourt, 2011). Therefore, all remaining subjects were exposed to a hybrid live-tutoring with passive song playback of that tutor that yielded more reliable tutor song imitation.

#### Live tutoring with audio playback – systemic subjects

Audio visual tutoring methods did not yield successful tutor imitations. Therefore, we opted for a tutoring paradigm that included a live-male tutor alongside passive audio playback as in London and Clayton (2008). Unlike some songbird species that can learn song from passive audio playbacks (e.g. Thorpe, 1958; Marler and Peters, 1988), zebra finches require either operantly-evoked playbacks or social instruction (Tchernichovski et al., 2001; Derégnaucourt, 2011; Deregnaucourt et al., 2013). We developed a tutor playback that combined passive audio playback alongside a live adult male. While operant playback has been used successfully to tutor zebra finches, we wanted to target the post-tutoring period with higher temporal precision. Operant training is pupil initiated and can span a long time period, whereas a controlled, timed playback allowed us to target the period immediately after training (i.e. the putative auditory memory consolidation period). To that end, we first identified a non-breeding adult male from our colony that was vocally active, and sang in the presence of an observer. The tutor was placed in a sound-attenuation chamber with an adult female and female-directed song was recorded, from which a 60-minute tutoring playback file was created. The same tutor playback and adult male was used for all systemically-treated subjects, as well as several of the microdialysis subjects (*n* = 22). After the original tutor perished, a new adult male was recruited, and a similar one-hour tutor playback file was created and presented to the remainder of subjects (*n* = 15).

The tutor playback file consisted of a 12-minute clip with 40 unique song bouts that was repeated five times. Each song bout contained 2 - 8 motifs, and included introductory notes. The 12-minute clip was assembled from 12 individual 1-minute blocks, where each block contained 30 seconds of song (4 – 5 song bouts per song period, each separated by 5 seconds of silence) followed by 30 seconds of silence. The final tutoring playback file was amplified to ∼70dB (A-weighted) and bandpass filtered at 0.3 – 15 kHz (Adobe Audition), and played through a portable speaker (Sony, model# SRS-TP1WHI) placed inside the sound-attenuation chamber.

The tutor was placed in an individual cage and kept in a sound-attenuation chamber with other adult zebra finches at least 24 hours before the day of tutoring. On the day of tutoring, an experimenter placed the tutor cage beside the pupil’s cage. After a 10-minute acclimation period without any song playback, the tutoring playback recording began. Immediately after the end of the tutor playback file, the tutor was removed from the pupil’s chamber.

### Bioacoustic analysis

#### Automated song analysis

Percent similarity, accuracy, and % sequence similarity was analyzed using SAP (Tchernichovski et al., 2000). Ten motifs of the tutor song were each compared to ten motifs of each pupil’s song from 130 dph using default settings for asymmetric mean values, yielding 100 comparisons per subject. Similar methods were used for measuring Weiner entropy (WE) and entropy variance (EV) across development in systemic subjects. As only half of the systemically-treated subjects produced song pre-tutoring (*n* = 3; 1 FAD subject and 2 saline subjects), we averaged pre-tutoring WE and EV across all subjects to compare with relative to 49 dph, which was the first day all subjects produced song.

#### Manual song similarity analysis

In addition to automated song similarity methods, we also measured the number of tutor syllables copied by each subject and the quality of each copy. Coded and randomized motifs were qualitatively analyzed on a syllable-by-syllable basis as being either ‘good’, ‘poor’, or ‘not available’ relative to the tutor song by three experimenters blind to treatment conditions and subject identification. We confirmed that raters agreed across multiple dimensions by performing inter-rater reliability measurements using an unweighted Fleiss’s Kappa. Raters were in excellent agreement in assessing syllable accuracy (K = 0.563, *p* < 0.001), assessing the syllables pupils were likely imitating (K = 0.657, *p* < 0.001), and on the total number of syllables copied from a tutor by a pupil (K = 0.455, *p* < 0.001). Moreover, raters’ intra-reliability was similarity high: raters agreed on 60.46% of syllable accuracy, 65.12% on pupil syllables that reflect the tutor syllable, and 58.14% on both the accuracy and imitated syllable in the pupil’s song. Further, raters’ similarity scores were well-matched to the SAP measurements: there was a significant positive correlation between all raters visual similarity scoring and SAP’s % similarity measurement (*r*(97) = 0.75, *p* < 0.001; ***Fig. 5C***).

#### Singing rate

An experimenter blind to treatment conditions measured the daily number of song bouts and their length for the entire pre-tutoring period (3 – 5 days pre-tutoring), tutoring period (10 days; tutor-off days), and every 5 days after the last day of tutoring until 130 dph (14 days). An individual song bout was defined as being at least 1 s in total duration and considered unique if 500 ms of silence elapsed between singing periods. Song bouts were analyzed for one 3-hour period per analyzed day (14:00 – 17:00). These methods were adapted from previous studies measuring song rate (Meitzen et al., 2007; Aronov et al., 2008; Meitzen et al., 2009; Alward et al., 2013).

#### Adult song plasticity

In a subset of birds (*n* = 23), we compared birds’ own song at 130dph (putative closure of the critical period for song learning) and song after being exposed to other adult male song (≥6 weeks post-130 dph return). We used simple qualitative measurements to assess whether song had changed (either ‘yes’ or ‘no’ based on visual comparisons of several song files from each time point) instead of more thorough bioacoustic analyses as treatment did not appear to affect the likelihood of changing adult song (see *Results*), which was the main question of the experiments.

### Behavior

#### Female two-choice song phonotaxis

Female songbirds use song to evaluate a potential mate (Zann, 1996; Tomaszycki and Adkins-Regan, 2005; Holveck and Riebel, 2007). Therefore, in addition to measuring song similarity, we also tested whether less subtle song features were affected by treatment by measuring song preference in adult female zebra finches. A 13” x 10” cage was placed in the center of a sound-attenuation chamber alongside speakers set on either side of it. Three ground-level perches were placed in the left- and right-most extreme side of the cage floor. A piece of cardboard cage matting was placed on the cage floor and divided into quarters with colored tape: left, left of middle, right of middle, and right. A non-breeding adult female zebra finch from our aviary (*N* = 12) was placed in the two-choice cage and isolated for ∼24 hours before the playback experiment began to increase salience of the future song playback. On Day 2, a 30 min song file was presented starting at ∼13:00. The song file consisted of a 2-minute clip repeated 15 times. The first minute of the 2-minute clip contained adult song solely from one FAD or saline bird, whereas the second minute of the 2-minute clip contained song from only one bird of the opposite treatment condition. Each 1-minute clip included 5 s of song, followed by 5 s of silence, which was repeated 4 times (40 s total), and followed by 20 s of silence (60 s total) played on one side of the speakers. The 1-minute clip of the second bird was broadcasted on the opposite speaker in a similar manner. The same 2-minute clip was repeated 15 times (30 mins total). On Day 3, a different playback file was played at a similar time (∼13:00) with new song stimuli played on opposite speakers compared to Day 2 to account for potential side-bias (e.g. if FAD song was broadcasted on the left speaker on Day 2, a new FAD song was broadcasted on the right speaker on Day 3). Females were returned to the aviary after the cessation of Day 3 playbacks. Birds were excluded from analysis if they spent the entire time in the middle/neutral zone (one bird was excluded from analysis from both days, and another bird was excluded from just one day of analysis). Total time spent near either the FAD or saline side was measured. Additionally, a FAD preference ratio was calculated similar to Remage-Healey et al. (2010):

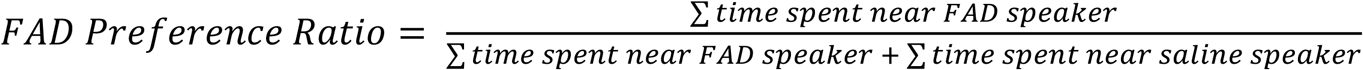

#### Microdialysis tutoring session behavior

Pupils who are more ‘attentive’ to the tutor during song learning sessions produce more similar copies of the tutor song in adulthood (Chen et al., 2016). As such, we explored whether treatment affected pupils’ behavior during tutoring sessions. Subjects were videotaped for 3 one-hour periods during each tutoring day, including: 1) the hour just prior to tutoring onset; 2) the tutoring period (∼70 mins; 10-minute acclimation period + 60 min audio playback); and 3) the hour immediately after tutor offset.

Three 10-minute clips per tutoring period for each subject were created for future behavioral scoring, including: 1) tutor acclimation period; 2) the beginning of tutor playback; and 3) 20 – 30 mins into the tutor playback period. Videos were scored for numerous behaviors by an experimenter blind to subjects’ treatment conditions using JWatcher (Blumstein and Daniel, 2007). Behaviors quantified included: events (eating; drinking; perch hops; grooming/preening; jumps; flights; feather ruffling; head scratching), and states (resting/sleeping; tutor zone; outside of tutor zone; not in view).

### Electrophysiology

#### Surgery

As others have reported (e.g. London and Clayton, 2008), guide cannulae implanted during development eventually dissociate from the skull, and the wound heals normally (see *Methods*), which allowed us to perform electrophysiology recordings from formerly dialyzed birds in adulthood. Surgical methods for the electrophysiology experiments were similar to previous procedures (Remage-Healey and Joshi, 2012; Ikeda et al., 2015; Vahaba et al., 2017; Krentzel et al., 2018), the main difference being a lack of an implanted microdialysis probe in the current experiment. At least six weeks following birds being returned to the aviary (median age on day of surgery = 227 dph, range = 158 – 526 dph), birds were recaptured, placed in a cage with a companion adult female, and song was recorded. On the day of the surgery, birds were initially food deprived for 30 minutes. Afterwards, an intramuscular injection of Equithesin was administered, and 20 minutes later, birds were swaddled in a Kim wipe, and placed atop a heating pad and secured via ear bars to a custom surgical stereotaxic apparatus (50° head angle). The bird’s beak was opened and placed in a beak holder. Once the bird was secured, head feathers were removed, and a 20 µL subcutaneous injection of 2% lidocaine was administered underneath the scalp, which was subsequently resected to expose the outer layer of skull. The MSB was then identified and used as our 0-point anatomical reference. A positioning needle was placed over the MSB, and adjusted to bilaterally mark NCM (rostral: -1.4 mm; lateral: ± 1.1 mm) and HVC (lateral: ± 2.40 mm). A piece of silver wire was inserted between the skull leaflets over the cerebellum to serve as a reference ground. A custom-fabricated metal head-post was then affixed above the beak and skull using dental cement and cyanoacrylate, followed by sealing the exposed scalp with cyanoacrylate. After surgery, birds were placed on heating pad within a recovery cage and provided with *ad libitum* food and water. Once birds awoke, they were returned to their sound-attenuation chamber in a separate cage from the companion female.

#### Anesthetized extracellular recordings

Extracellular, multiunit electrophysiological recordings were obtained from NCM and HVC in anesthetized subjects (*n* = 21 birds [aCSF = 8 birds; FAD = 8 birds; cannula = 5 birds]; single-units x treatment x region: aCSF = 20 HVC units; 49 NCM units; FAD = 31 HVC units; 48 NCM units; cannula = 14 HVC units; 18 NCM units) using Spike2 (version 7.04, Cambridge Electronic Design) at a sampling rate of 16.67 kHZ, bandpass-filtered at 0.3 – 5 kHz. On the day of the experiment, birds were starved for 30 mins, followed by three intramuscular injections of 20% urethane on alternating sides of pectoral muscle (∼100 µL total; ∼33 µL per injection). Injections were administered every 45 minutes. Once anesthetized, birds were brought up to the recording room, wrapped in a Kim wipe, placed on a heating pad, and affixed to a custom head-post stereotaxic apparatus. The outer- and inner-leaflet of skull and dura was then exposed over the HVC and NCM of one hemisphere. A drop of silicone oil was placed over the exposed brain to prevent the tissue from drying out. Individual carbon-fiber electrodes (CarboStar-1; Kation) were advanced into the proximate region of NCM and HVC based on: 1) anatomical location (∼0.80 – 1.40 mm ventral; and ∼0.50 mm ventral, respectively); and 2) characteristic spontaneous- and stimulus-evoked firing rates. In anesthetized adult songbirds, HVC preferentially responds to playbacks of birds’ own song (BOS) (Margoliash, 1983, 1986). As such, we played BOS along with other songs in our search stimuli set (see below) and used a combination of characteristic spontaneous activity and neural responses to BOS as an indication of placement within HVC. After a completed playback trial, electrodes were advanced 100 – 150 µm dorsal/ventral along the same track, and, if the region-specific characteristic firing persisted, a new recording was obtained. Once a track was past anatomical limits and/or ceased to display characteristic firing patterns, an electrolytic lesion presented at the most recent site for future anatomical confirmation. After one hemisphere was complete, the contralateral hemisphere was exposed and recorded. At the end of the experiment, birds were rapidly decapitated, and their brains were extracted and placed in a 20% sucrose-formalin solution for future sectioning and histology. In addition to recording from successful subjects (i.e. aCSF and FAD treated subjects), we also recorded from subjects that whose microdialysis cannulae became non-functional during the tutoring experiment. We present these data as a visual comparison as surgery controls (noted as ‘Cannula’ subjects) but due to the variability for microdialysis failure in these subjects, we omitted them from our statistical model.

#### Auditory stimuli and playback

All stimuli were adjusted to ∼70 dB (A-weighted) and bandpass filtered to 0.3 – 15 kHz (Adobe Audition). Two sets of stimuli were used during the recordings. A *search* set was composed of two unique conspecific songs (i.e. zebra finch; CON), birds’ own song (BOS), reverse BOS (REV-BOS), and white noise (WN). The *experimental* set was composed of two novel CONs, BOS, REV-BOS, tutor’s song (TUT), reverse TUT (REV-TUT), and WN. Search stimuli were presented manually by the experimenter to confirm putative NCM and HVC sites, whereas the experimental set were played automatically and randomized via a custom written script in Spike. For experimental playbacks, each stimulus was pseudorandomly played once per block, with a total of 20 blocks being presented for each playback period. Stimuli were separated by a 10 s inter-stimulus interval ± 0 - 2 s of random time.

#### Single-unit spike sorting

Individual single units were sorted using default parameters in Spike2 (v.7.04, Cambridge Electronic Design; as in Vahaba et al., 2017). Units were retained for analysis if they: 1) were distinctly clustered in a principal component analysis space (apart from noise and other units); 2) had an interspike interval of > 1 ms; and 3) were auditory responsive by visual inspection of the peristimulus time histogram and raster plots.

#### Data analysis

Single-unit electrophysiology recordings were analyzed using similar methods as in Vahaba et al. (2017). Briefly, spontaneous firing rates were defined as the total number of waveform events (spikes) occurring in a 2-second period prior to the onset of an auditory stimulus, whereas stimulus-evoked firing was defined as the number of spikes during a 2-second window starting at the onset of an auditory stimulus. The total number of spikes per stimulus were divided by the number of stimulus iterations to yield firing rates (Hz). Firing rates were also z-transformed to normalize data and account for variability across subjects and units using the following equation:

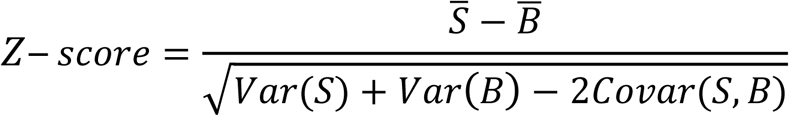

Where *S* and *B* represents the number of, respectively; *S̅* and *B̅* represent the mean number of stimulus and spontaneous spikes for a given stimulus.

We also analyzed stimulus selectivity using d prime (d’; Green and Swets, 1966), a psychophysics metric of discriminability used for assessing neural responses to a given stimulus relative to a different stimulus (e.g. Bauer et al., 2008; Remage-Healey and Joshi, 2012; Moseley et al., 2017), using the following equation:

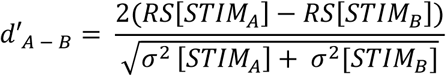

Where *RS* is the response strength (mean stimulus-evoked firing rate subtracted from the mean spontaneous firing rate), STIM_A_ represents the focal stimulus of interest, STIM_B_ represents the relative stimulus to compare other stimuli to, and *σ*^2^ is the RS variance for a given stimulus. WN was used as the comparison stimulus for NCM recordings, and CON1 for HVC recordings (see *Results*).

#### Adult Habituation Experiment

##### Subjects

A separate set of otherwise untreated adult male zebra finches (*n* = 22) were removed from our single-sex aviary (median age on day of electrophysiology recording = 274 dph; all males at least 120 dph) and placed in a cage within a sound-attenuation chambers alongside an adult companion female while song was recorded using Sound Analysis Pro (Tchernichovski et al., 2000). Birds were kept in the same cage until the day of the surgery which typically occurred after 3 days of isolation (mode = 3 days isolation pre-surgery; range = 0 – 6 days).

##### Surgery

The surgery methods used for this experiment were nearly identical to the one above. The main difference was that the skull was exposed solely over both hemispheres of NCM. After bilateral marking of NCM (coordinates = rostral: -1.20 mm; lateral = +/- 1.10 mm), the outer and inner leaflets of skull were carefully removed, leaving the dura intact as much as possible. Following silver wire implantation, a silicone dural sealant (Kwik-Sil, World Precision Instruments [WPI]) was placed over the exposed skull.

##### Auditory Training & Drug Administration

Awake birds were placed in a custom-fabricated restraint tube and brought into the recording room. After being secured to the head-post stereotaxic apparatus, 200 iterations of a single adult male zebra finch song (two motifs within one song bout, including intro notes) was presented (TRAIN) with a 12 s ISI. Training lasted 46 minutes in total. Immediately after the last TRAIN iteration, ∼100 nL of either artificial cerebrospinal fluid (aCSF) or 100 µM FAD in aCSF were locally administered via pre-loaded glass micropipettes broken back to ∼24 µm internal diameter, which were left in place for >2 minutes following injection to prevent dispersal. This volume has been successfully used in previous studies and appears to disperse across the extent of NCM (Tremere et al., 2009; Remage-Healey and Joshi, 2012). Pipettes were successively descended ventral 1.10 mm in NCM, followed by pressure-injections (Pneumatic PicoPump, PV830; World Precision Instruments). Following drug treatments, the exposure was sealed with a lower tear-strength silicone adhesive (Kwik Cast), cured, and then the bird was returned to his cage.

##### Electrophysiology

Awake, restrained birds were brought back to the recording room for electrophysiology recordings 6 or 20 hours after training. Birds were non-anesthetized as habituation is not typically observed in anesthetized songbirds (Remage-Healey et al., 2010), but see (Ono et al., 2016). Parylene-coated tungsten electrodes (0.5 or 2MΩ; A-M Systems) were descended bilaterally into the approximate drug injection region from *Training*. Recordings were amplified using an A-M system amplifier and obtained through a connected 1401 board and Spike2 (CED). A set of stimuli were first presented to the bird *search stimuli* to confirm the recording site displayed NCM characteristic-like auditory responses. After site confirmation, experimental stimuli were presented to the bird while neural activity was continuously recorded. Each recording site was electrolytically lesioned following playback. Recording sites/exposures were once again covered with silicone adhesive, and birds were either sacrificed via rapid decapitation immediately after recordings (*n* = 8) or 2-3 days later (*n* = 15) to allow for lesion sites to become more pronounced and readily observable in sectioned tissue (e.g. allow time for gliosis). Extracted brains were placed in 20% sucrose-formalin for attempted future sectioning and histological verification of recording and drug site via Nissl stain.

##### Auditory Stimuli & Playback

All auditory stimuli were presented at ∼70 dB. Search stimuli consisted of a unique set of non-familiar conspecific song not used in the experimental stimuli set. Experimental playback stimuli presented during neural recordings included the trained conspecific song (TRAIN) and its reverse (REV-TRAIN), three novel conspecifics (CON1, CON2, CON3) and one reversed (REV-CON3), bird’s own song (BOS) and its reverse (REV-BOS), and white noise (WN). To ensure birds were unfamiliar with the song presented, several stimuli were graciously adapted from an online zebra finch song repository (http://people.bu.edu/timothyg/song_website/). We also used two songs from birds in our own breeding colony as they had been removed long before the experiment began. Birds were presented with 25 consecutive iterations of each experimental stimulus with a 12 second ISI in blocks (e.g. 25 CON1 playbacks, then 25 CON2 playbacks, then 25 WN playbacks, etc.), as in previous experiments (e.g. Yoder et al., 2012).

##### Analysis

Analyses were inspired from previous studies with minor changes (e.g. Yoder et al., 2012). Briefly, multi-unit recordings were analyzed root mean squared (RMS) in Spike2 for the stimulus and baseline period. The stimulus period included the entire duration of playback stimulus + 100 ms after offset, whereas the baseline period was defined as a 500 ms period preceding stimulus onset. Mean baseline RMS was derived across the entire recording period, whereas mean stimulus RMS was calculated for each individual stimulus. Data were filtered on a per trial (i.e. stimulus repetition) basis. First, any trial exceeding two-times the mean RMS for either baseline or stimulus RMS (separately) was excluded. After, any trial above/below 2.5 standard deviations was then excluded for both stimulus and baseline RMS. Finally, a grand mean baseline RMS (derived from the entire recording period; across stimuli) was subtracted from stimulus RMS values, yielding an adjusted RMS. Slope was derived from trials 1 – 25 using the *lm*() function via the *stats* package in R.

### Statistical analysis

All statistical analyses were performed using R (R Core Team, 2018) via RStudio (RStudio Team, 2016) using several packages, including: *tidyverse*; *plyr*; *sciplot*; *irr*; *corrplot*; *data.table*; and *Hmisc*. Histology data (% DAPI; cell density) were analyzed using a two-way ANOVA (NCM subregion X phase). Singing rates were analyzed using a two-way ANOVA (treatment X time of day). Pearson’s correlation was used to analyze changes in Weiner entropy relative to eventual song similarity at 130dph. One-way ANOVAs were employed to assess systemic treatments effect on song learning outcomes (separate analyses for per cent similarity, sequential similarity, and accuracy). Female phonotaxis data were analyzed using two-way ANOVAs (treatment X trial day). For microdialyzed subjects, automated and manual song similarity analyses were analyzed using a two-way ANOVA (treatment X hemisphere). Inter-rater reliability for manual song similarity scoring was analyzed using an unweighted Fleiss’s kappa. The comparison between automated (SAP) and manual (visual) song similarity was measured using Pearson’s correlation. Tutoring behavior was analyzed using a mixed-effects ANOVA (tutoring day [within] X treatment [between]), and a correlation matrix adjusted for multiple comparisons (adjusted α = 0.00048). For behavioral analyses, we restricted our data to the first 10 minutes of tutoring for only days 1 and 2 to be consistent as some subjects received three days of tutoring. A chi-squared was used to compare distributions of adult song plasticity across treatment. For electrophysiology measurements, a three-way ANOVA was employed (treatment X recording hemisphere X stimulus). Finally, for adult habituation neural recordings, adaption slopes were compared using two-way ANOVAs (treatment type [aCSF/nothing vs. FAD] X stimulus type [familiar vs. trained]). All *post hoc* comparisons were performed using Tukey’s honestly significant difference (HSD) test, and were corrected for multiple comparisons. *P*-values < 0.05 were considered significant. Data from ‘cannula’ subjects were omitted from any statistical model and are plotted throughout the manuscript as a visual comparison (see *Results*).

## RESULTS

### Cell density is region- and age-dependent in developing auditory forebrain while aromatase and parvalbumin expression are unchanging

We first sought to confirm the presence of aromatase in NCM across development. While previous studies have characterized aromatase expression in the brains of developing songbirds, both directly (protein: Saldanha et al., 2000) and indirectly (Vockel et al., 1988; Jacobs et al., 1999; Saldanha et al., 1999; Chao et al., 2015), information on aromatase protein expression specifically within NCM between the sensory and sensorimotor phase of the song learning period has not been assessed, to our knowledge. In addition to aromatase, we were also curious as to whether transitions between learning phases were associated with differences in expression of the calcium buffering-protein parvalbumin. Parvalbumin is a marker for a unique subpopulation of inhibitory interneurons (Tremblay et al., 2016), is co-localized with aromatase in NCM (Ikeda et al., 2017), and its presence often denotes changes in critical period plasticity within mammalian visual cortex (Hensch, 2005), as well as songbird song circuits (Balmer et al., 2009). We focused solely on males as they were the sex of interest for subsequent song and physiology experiments in this study. Although we collected both hemispheres of NCM for this experiment, we excluded hemisphere as a factor in our statistical model to support sufficient statistical power for our comparisons. Qualitatively, we found similar expression of aromatase and parvalbumin across both hemispheres of sensory- and sensorimotor-aged subjects (see ***Tables 1 & 2***).

**Table 1.** Density measures for antibody staining in developing NCM. Values represent mean cell density (mm^3^) +/- the standard error of the mean.

We divided our subjects into two age groups reflecting the two different developmental song learning phases: sensory- and sensorimotor-aged (20-34 and 65-71 dph, respectively, (Vahaba et al., 2017)). Overall, density measures revealed comparable aromatase, parvalbumin, and aromatase-parvalbumin co-expression in both dorsal and ventral NCM across development (aromatase: *F*_(1, 31)_ = 2.458, *p* = 0.127; parvalbumin: *F*_(1, 31)_ = 0.035, *p* = 0.854; aromatase-parvalbumin: *F*_(1, 31)_ = 0.003, *p* = 0.957), age (aromatase: *F*_(1, 31)_ = 2.218, *p* = 0.147; parvalbumin: *F*_(1, 31)_ = 0.277, *p* = 0.602; aromatase-parvalbumin: *F*_(1,31)_ = 0.339, *p* = 0.565), without any significant interactions between age and region (aromatase: *F*_(1, 31)_ = 0.048; parvalbumin: *F*_(1, 31)_ = 0.751; aromatase-parvalbumin: *F*_(1, 31)_ = 0.757; *p* > 0.3 for all tests; ***Table 1***).

Interestingly, we observed a significantly higher DAPI expression in dorsal NCM compared to ventral NCM (*F*_(1, 31)_ = 8.128, *p* = 0.008), as well as higher DAPI expression in sensory-aged animals compared to sensorimotor-aged subjects (*F*_(1, 31)_ = 6.291, *p* = 0.018; ***Fig. 2C,D***). No significant interactions emerged between region and age (*F*_(1, 31)_ = 0.587, *p* = 0.449). There were no differences between age and NCM subregion when we normalized the markers of interest (aromatase and parvalbumin) to the relative amount of DAPI to account for subject and image variability (***Fig. 2A,B***; see ***Tables 1 & 2*** for all descriptive data for density and % of DAPI measurements). Overall, these findings confirm that aromatase and parvalbumin are present in the developing auditory forebrain, and that NCM appears to undergo cellular pruning as birds develop while maintaining subregion differences in cell density.

**Fig. 2.**
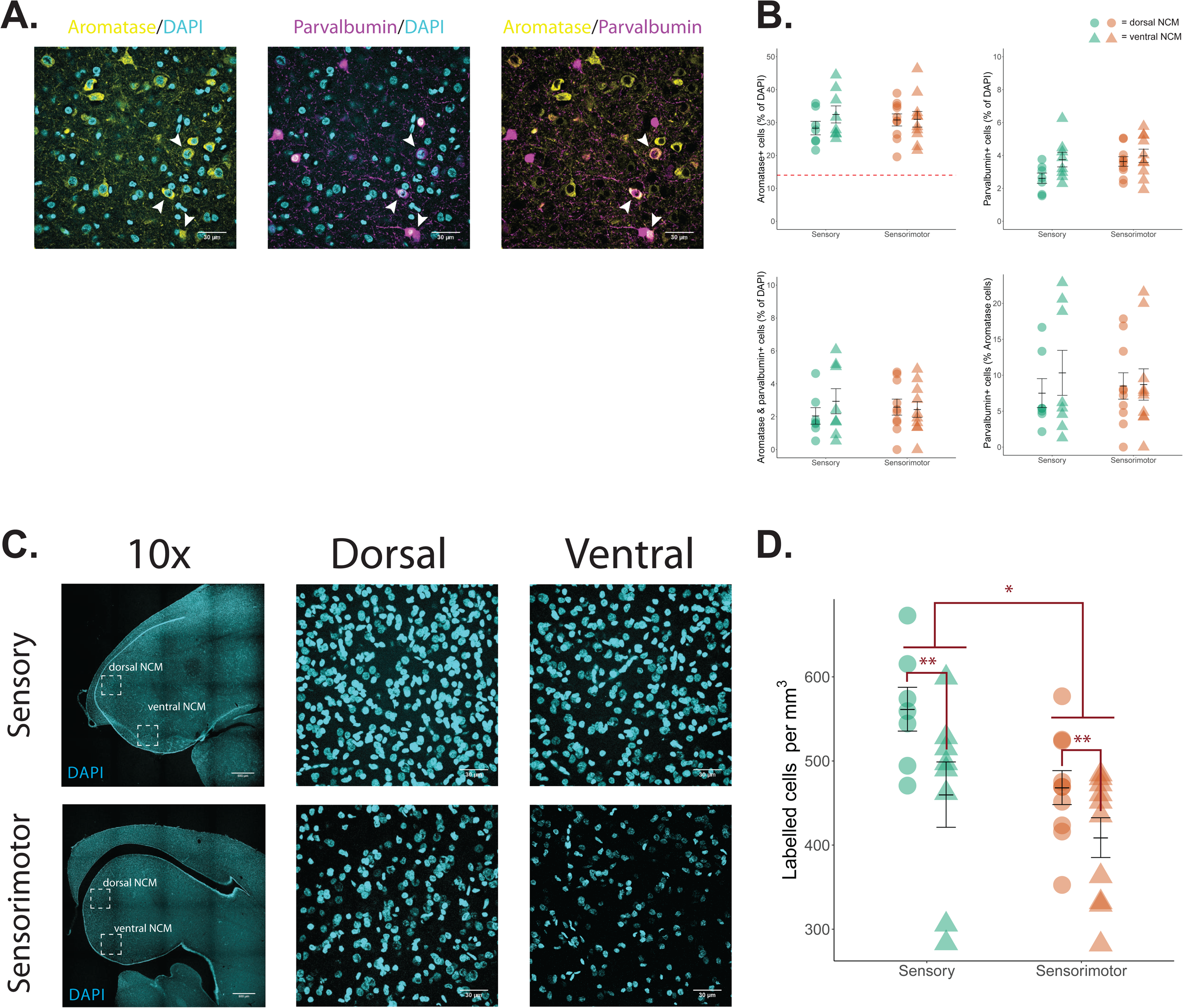
Changes in neuronal density and aromatase and parvalbumin expression in NCM across development. **A**, Aromatase, parvalbumin, aromatase parvalbumin co-expression, respectively, from an exemplar sensory-aged male bird (26 dph; right hemisphere; ventral NCM). Pseudo-colored: yellow, aromatase; cyan, DAPI; magenta, parvalbumin. Each image from a single slice of a z-stack taken at 60x magnification. Scale bar = 30 µm. White arrowheads indicate aromatase and parvalbumin co-expression. **B**, Expression of aromatase, parvalbumin, and aromatase/parvalbumin co-expression, respectively, relative to the expression of DAPI (%), and parvalbumin co-expression relative to total aromatase expression (%). Overall, there are no significant differences in expression by age or NCM subregion. Circles = dorsal NCM; triangles = ventral NCM; green = sensory-aged birds; orange = sensorimotor-aged birds. **C**, DAPI expression across development; *top row*: sensory-aged bird (25 dph; right NCM); *bottom row*: sensorimotor-aged bird (71 dph; right NCM). 10x images taken from a 4 x 4 stitched image. Dorsal and ventral NCM images taken from a z-project max intensity 60x image. **D**, Cell density (DAPI expression) by region and age. Dorsal NCM shows higher cell density than ventral NCM. Similarly, sensory-aged birds have higher overall cell density across subregions compared to sensorimotor-aged subjects. * = *p* < 0.05; ** = *p* < 0.001.

**Table 2.** Protein expression relative to cell density (% DAPI). Values represent mean number of immunopositive-cells relative to the number of DAPI+ neurons (% DAPI) +/- the standard error of the mean.

### Song learning is unaffected by global estrogen synthesis inhibition during development

#### Systemic administration

Birds in this experiment received an oral administration of either FAD or saline every other day for 20 days immediately following tutoring. Initially, we measured singing rates of systemically-treated animals before (<40 dph) and during the tutoring period (40 – 60 dph) as global inhibition of estrogen synthesis in adult songbirds can reduce song production (Alward et al., 2016). Pre-tutoring, birds sang at comparable rates independent of the time of day or future treatment group (treatment: *F*_(1, 13)_ = 2.466, *p* = 0.140; time of day: *F*_(1, 13)_ = 1.797, *p* = 0.203; treatment * time of day: *F*_(1, 13)_ = 0.719, *p* = 0.412; ***Fig. 3A,B***). However, during the tutoring period, FAD treatment significantly suppressed singing rates (FAD = 63.8 ± 13.6 bouts; saline = 116.0 ± 14.4 bouts; *F*_(1, 103)_ = 6.623, *p* = 0.012; Tukey’s HSD: *p* = 0.012) independent of time of day (*F*_(1, 103)_ = 0. 222, *p* = 0. 639) or an interaction between time of day and treatment (*F*_(1, 103)_ = 1.882, *p* = 0.173; ***Fig. 3A,B***). Interestingly, while initial song production was reduced during development, eventual song similarity at 130 dph (one-way ANOVA (treatment); *F*_(1, 4)_ = 0.064), accuracy (*F*_(1, 4)_ = 0.021), and sequential similarity (*F*_(1, 4)_ = 0.095) were statistically similar when both FAD and saline subjects reached adulthood (*p* > 0.77; ***Fig. 3D & Table 3***). Additionally, while FAD birds appeared to exhibit a lower tutor song similarity score early in development (49 dph), there was no effect of treatment (*F*_(1, 4)_ = 0.427, *p* = 0.549), nor an interaction of treatment with age (F(4, 16) = 0.569, *p* = 0.689). There was, however, a significant increase in song similarity as birds reached adulthood (age: *F*_(4, 16)_ = 5.528, *p* = 0.005; *post-hocs*: *p* < 0.05 for 49 dph vs. 86 & 130 dph; all other age comparisons non-significant, *p* > 0.06; ***Fig. 3C***). Together, these data show that systemic estrogen synthesis can support song production during the juvenile learning period but it does not impact eventual tutor song imitation.

**Fig. 3.**
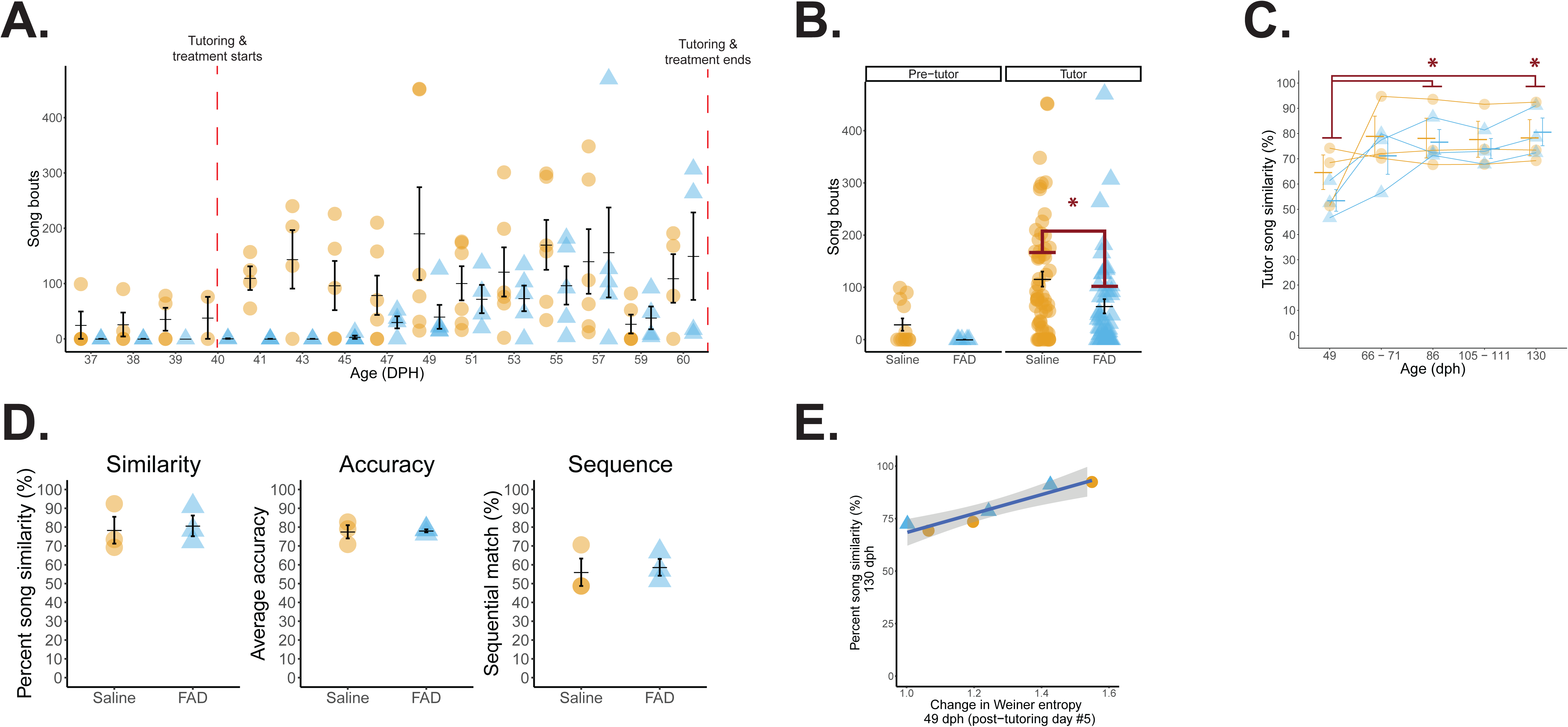
Systemic estrogen synthesis inhibition suppresses song production without impacting tutor song copying. **A**, Daily number of song bouts before and across the tutoring/treatment period. **B**, Birds sing at similar rates before treatment/tutoring; however, systemic FAD treatment reduces song production (*p* = 0.012). Circles/orange = saline-treated birds (*n* = 3); triangles/blue = FAD birds (*n* = 3). **C**, Song similarity is lowest at 49 dph despite treatment (effect of age: *p* = 0.005; * is relative to 49 dph). **D**, At 130 dph, tutor song similarity, accuracy, and sequence similarity, respectively, are all similar across treatments. **E,** Change in Weiner entropy at 49 dph (post-tutoring day #5) predicts eventual percent song similarity to the tutor at 130 dph, independent of treatment (*r*2 = 0.903*; p* = 0.004). * = *p* < 0.05.

**Table 3.** Automated song similarity measurements. Values represent mean +/- the standard error of the mean for each song similarity metric.

Developmental changes (relative to pre-tutoring values) in Weiner entropy (WE) and entropy variance (EV) during tutoring predict adult tutor song fidelity (Deshpande et al., 2014). Independent of treatment, we tested this relationship for birds in the present experiment to assess whether they developed along a ‘typical’ song learning trajectory. In agreement with the previous report, we found a strong, significant positive correlation between change in WE at 49 dph and percent song similarity in adulthood (130dph); *r*(4) = -0.951, *p* = 0.004, as well as a similar significant correlation when we considered entropy variance instead of WE (*r*(4) = 0.863, *p* = 0.027; ***Fig. 3E***). Therefore, while systemic FAD treatment did not impact song learning, developing song was predictive of eventual similarity, indicating that our daily treatment regimen did not impair a ‘normal’ song learning trajectory.

#### Female phonotaxis behavior

While song similarity data can provide information on how well a bird imitates a model song, there are likely subtle song features that are affected by early-life manipulations that may not be captured by automated analyses. As adult female zebra finches use courtship song to evaluate potential life-long mates (Zann, 1996), we asked whether a females’ song preference was impacted by a males’ drug treatment during development. We found a significant interaction between treatment and trial day (*F*_(1, 17)_ = 7.30, *p* = 0.151). Follow-up analyses revealed that on the first day of phonotaxis, females spent more time near the speaker broadcasting a FAD-treated birds’ song (*p* = 0.015), whereas on the second day there was nonsignificant tendency for preferring control birds’ song (*p* = 0.059; ***Fig. 4A***). We also evaluated a ‘FAD preference ratio’ for day 1 vs. day 2. Visually, it appears that females initially prefer FAD song and then ‘switch’ preferences on day 2, but this was not statistically significant (*F*_(1, 8)_ = 2.958, *p* = 0.124; ***Fig. 4B***). We also confirmed that there was no overall side bias (*p* = 0.0989) nor inherent preference for a specific male’s song independent of treatment (*p =* 0.557). Thus, systemic estrogen synthesis blockade during development did not negatively impact song features important for eventual female song preference.

**Fig. 4.**
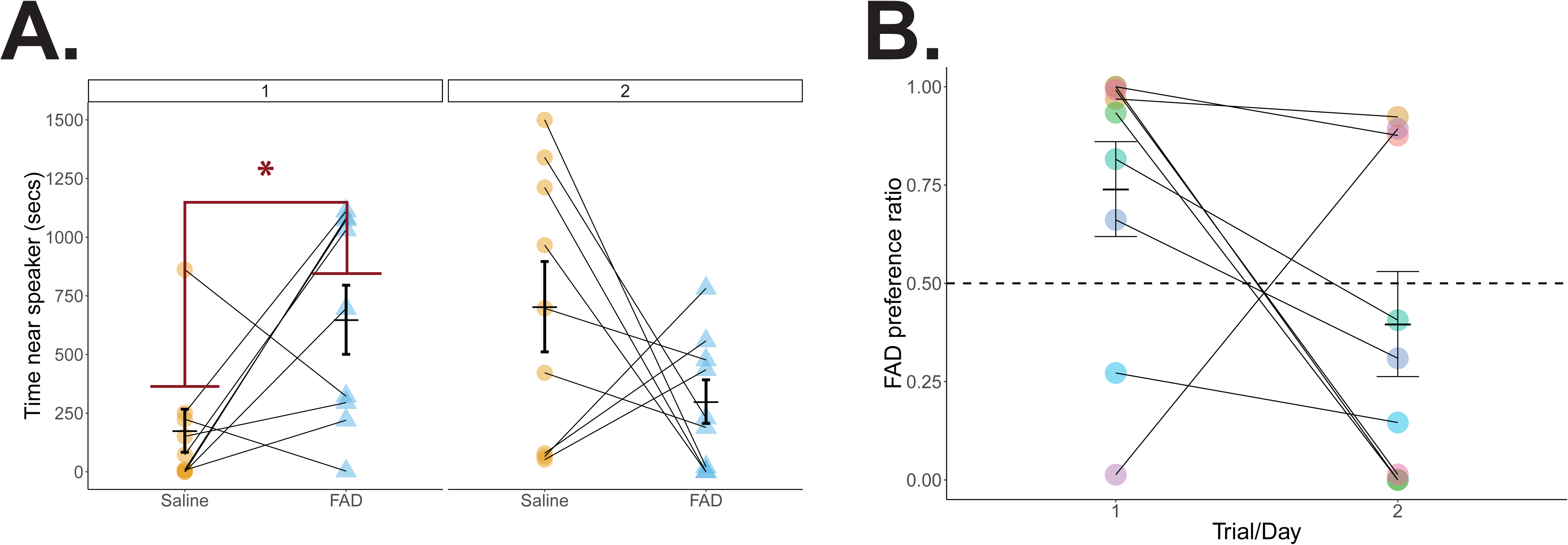
Female songbirds temporarily prefer E2-suppressed adult song. **A**, Adult female songbirds spend more time near a speaker broadcasting adult song from systemic E2 suppressed males on day 1, but not 2, in a two-day phonotaxis experiment (*p* = 0.015). **B**, Preference ratios for FAD song relative to control song is similar across days. * = *p* < 0.05.

**Fig. 5.**
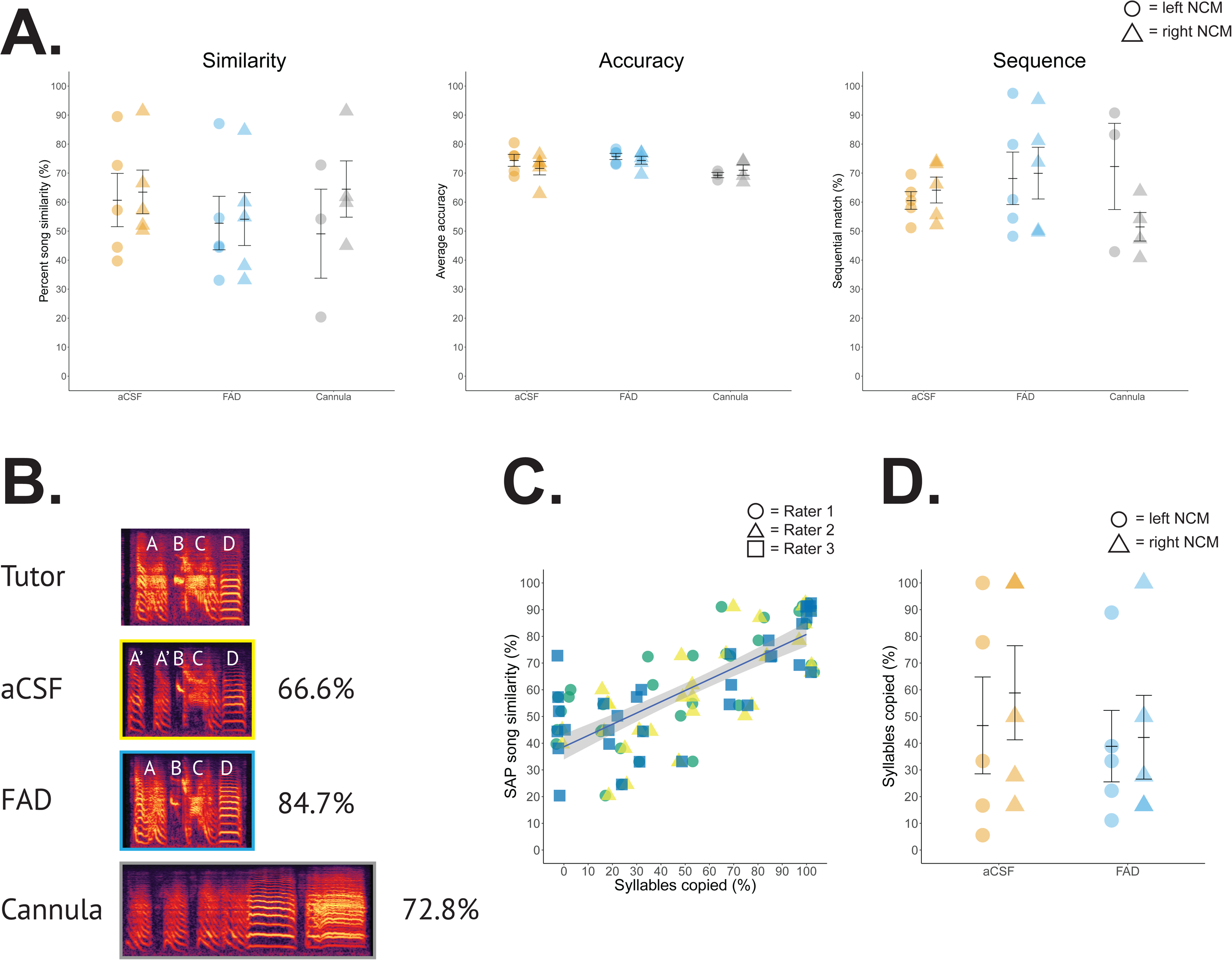
Song copying is unaffected by central estrogen production inhibition via *in vivo* microdialysis. **A**, 130 dph song similarity, accuracy, and sequence similarity, respectively, are all comparable across aCSF- and FAD-treated birds. Cannula ‘surgery controls’ are graphed for visual comparison. Orange = aCSF; blue = FAD; grey = cannula; circle = left NCM; triangle = right NCM. **B**, Example song spectrograms and their average song similarity % relative to tutor. Letters denote syllables; A’ = partial syllable derived from A. Note the seemingly high similarity of both the aCSF and FAD motif, yet divergent song similarity scores (aCSF bird = right NCM; FAD bird = right NCM; similarity score is averaged across 100 motif comparisons, see *Methods*). **C**, Manual song similarity measurements are strongly correlated with automated methods; color/shape denotes unique rater (*r*^2^ = 0.563, *p* < 0.001); jitter added to reveal overlap. **D**, As with automated methods, manual song similarity scoring reveals comparable tutor song copying across treatments.

### Song learning is unaffected by inhibition of local estrogen synthesis in NCM during development

#### In vivo microdialysis with social + playback tutoring

Systemic treatments yielded no effect of aromatase blockade, but leaves open the possibility that temporally-precise, site-directed manipulations within NCM could impact auditory memorization. However, as with systemically-administered subjects, central unilateral FAD treatment did not modify eventual tutor imitation, nor did the cannulated hemisphere or interaction between treatment and hemisphere affect percent similarity (*F*_(1, 16)_, treatment = 0.965; hemisphere = 0.056; treatment * hemisphere = 0.007; *p* > 0.340), accuracy (*F*_(1, 16)_, treatment = 1.325; hemisphere = 1.277; treatment * hemisphere = 0.157; *p* > 0.266), or sequence similarity (*F*_(1, 16)_, treatment = 0.950; hemisphere = 0.153; treatment * hemisphere = 0.017; *p* > 0.343; ***Fig. 5A & Table 3***). Therefore, contrary to our original prediction, unilateral central estrogen synthesis blockade in NCM did not impair tutor song memorization and eventual imitation.

#### Manual song similarity quantification

Whole motif similarity measurements via SAP is the conventional method to objectively analyze tutor similarity for zebra finches (Tchernichovski et al., 2000). Inspection of spectrograms suggested that SAP similarity measurements were not capturing the full extent of tutor song similarity (***Fig. 5B***: high % SAP song similarity for *Cannula* subject, but visually and acoustically dissimilar; opposite issue with *aCSF* subject). To address this, we employed visual song similarity measures in the spirit of early songbird bioacoustic research studies that relied solely on visual spectrographic assessment (Borror and Reese, 1953; Thorpe, 1954; Eales, 1985). In accordance with this match between SAP and when visual scoring methods, there were no significant effects for visually-scored song similarity (average percent copied) by cannulated hemisphere (*F*_(1, 16)_ = 0.227, *p* = 0.640), treatment (*F*_(1, 16)_ = 0.561, *p* = 0.465), nor an interaction between either factor (*F*_(1, 16)_ = 0.074, *p* = 0.789; ***Fig. 5D***). Therefore, irrespective of bioacoustic assessment, unilateral blockade of neuroestrogen production in the auditory forebrain during and immediately after song learning did not impair auditory memorization of the tutor song.

#### Tutoring behavior

Attention plays a critical role for vocal learning early in development (e.g. Chen et al., 2016). Since estrogens can modulate attention in rodents (see references in Sommer et al., 2018) we explored whether FAD treatment impaired measures of attention during tutoring sessions in a subset of subjects (FAD *n* = 9; aCSF *n* = 9). Overall, we found no effect of treatment on the amount of time pupils spent near the tutor (‘tutor zone’; a proxy for tutor attention) on either tutoring day (*F*_(1, 14)_; *p* > 0.190 for main effects and interaction; ***Fig. 6A***). The time spent near the tutor is one obvious behavior to explore with clear predictions about its impact on eventual song copying. However, as there are not many quantitative data to our knowledge on pupil behavior during tutoring, we also explored whether the other behaviors we scored might also be predictive of future tutor song similarity. We generated a correlogram that included all tutor session behaviors, as well as song similarity measurements (both visual and SAP derived). Overall, there were few significant correlations of interest pertaining to song similarity and behavior that emerged (***Fig. 6B***).

**Fig. 6.**
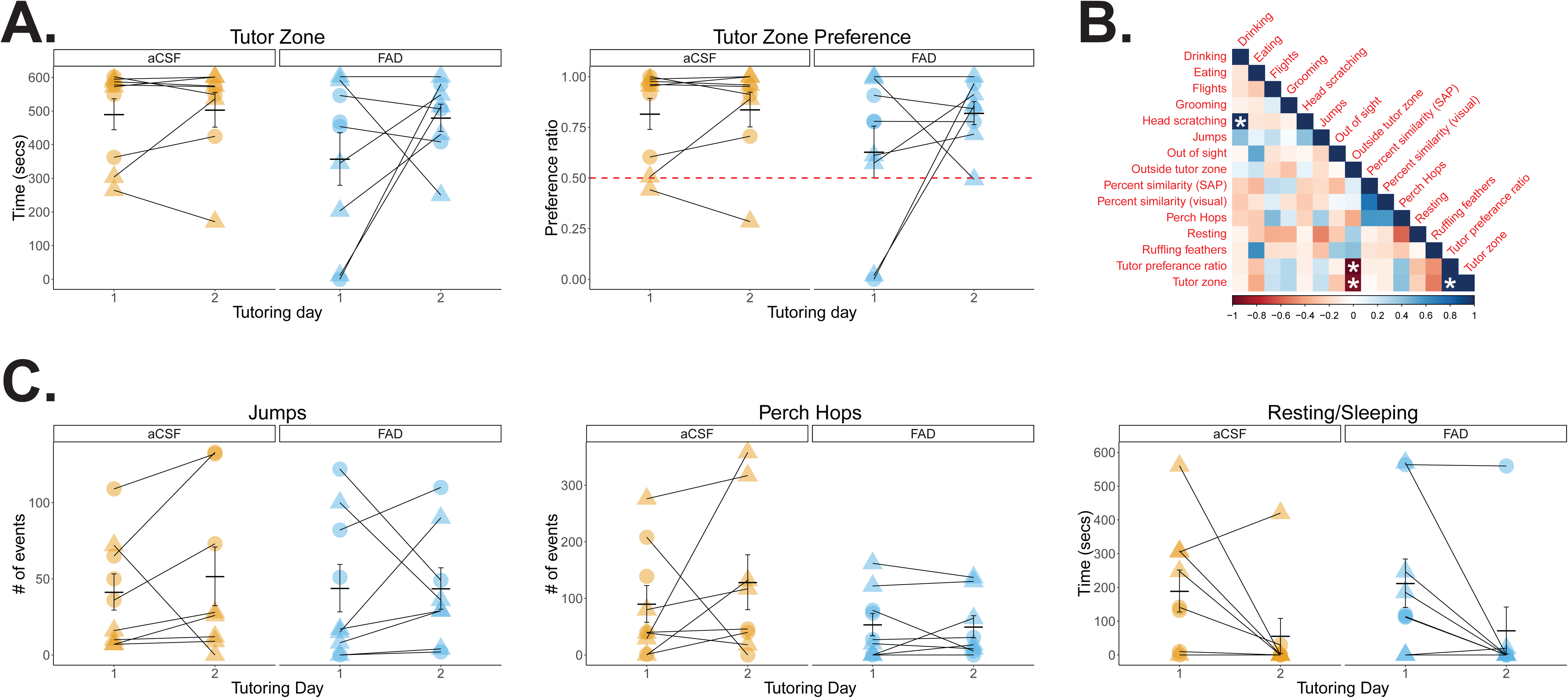
Juvenile male songbirds are similarly attentive to the tutor during microdialysis. **A,** The time a bird spent near a live adult male tutor during *in vivo* microdialysis is similar across treatments, targeted hemispheres, and tutoring day. Behavior presented is from the first 10 minutes of song playback alongside live male presentation (see *Methods*). Orange = aCSF; blue = FAD; circle = left NCM; triangle = right NCM. Tutor preference ratios are similar across treatments. **B**, Correlogram of tutoring behavior and song similarity measurements reveal significant correlations (more time spent near the tutor negatively associated with time spent away from the tutor; tutor zone time positively correlated with tutor preference ration), and novel findings (positive correlation of head scratching and drinking); *p* < 0.0005 (adjusted α; Bonferroni correction). Behavior data presented is from the first 10 minutes of tutor playback across days 1 and 2 of tutoring. **C**, Motor activity is statistically similar across treatment and tutoring days.

Another possibility is that FAD treatment may impair locomotion. We explored whether two common motor behaviors (jumping and perch hopping), as well as time spent resting/sleeping were affected by pharmacological exposure. Overall, neither treatment nor tutoring day affected jumping or perch hops (*F*_(1, 14)_; *p* > 0.158 for main effects and interaction; ***Fig. 6C***); however, birds spent more time resting irrespective of treatment on the second day of tutoring, suggesting that the novelty of an adult male wanes after the first session (F(1, 14) = 7.938, *p* = 0.0137; all other analyses *p* > 0.808; ***Fig. 6C***). These results suggest that as with song similarity, behavior during a social learning session is similarly unaffected by unilateral central neuroestrogen synthesis blockade.

#### Song changes after exposure to adult male conspecifics

We noticed highly aberrant song types in several formerly microdialyzed subjects independent of treatment at 131 dph (*X*2 (*N* = 23) = 1.189, *p* = 0.552), which is well beyond the putative ‘closing’ of the critical period for song learning and song should be highly stable (***Supp. Fig. 1***). Aberrant songs were always highly variable (i.e. not crystallized/stereotyped) at 130 dph and eventually reverted to higher stereotypy after being exposed to other adult male birds, and typically involved dropping and/or adding new syllables (6/8 subjects added, dropped, or modified syllables). These results suggest that, in addition to age, experience gates the song learning critical period closure, which has been described in other studies on lab-reared tutored and isolate zebra finches (Eales, 1985; Morrison and Nottebohm, 1993; Jones et al., 1996; Deregnaucourt et al., 2013).

### Neuroestrogen suppression in development leads to enhanced neural representations of birds’ own song and tutor song in HVC of adults

In a subset of formerly microdialyzed birds (21 out of 28 birds), we obtained neural recordings from two brain regions associated with song learning and tutor memory representation: NCM and HVC. Recordings were obtained from both the contralateral and ipsilateral hemisphere relative to the site of microdialysis cannulation (i.e. contralateral = recording from non-dialyzed hemisphere; ipsilateral = recording from dialyzed hemisphere).

#### NCM

We first explored whether treatment impacted NCM firing properties. Spontaneous firing rates were unaffected by recording hemisphere, treatment, and there was no interaction between the two factors (*F*_(1, 93)_ = 0.238, 0.003, and 0.779, respectively; *p* > 0.60; ***Fig. 7A***). Contrary to spontaneous firing, stimulus-evoked firing was significantly affected by a recording hemisphere x treatment interaction (*F*_(1, 651)_ = 7.938, *p* = 0.005) as well as there being a main effect for treatment (*F*_(1,6)_ = 4.334, *p* = 0.038) and stimulus (*F*_(6, 651)_ = 7.670, *p* < 0.001). Follow-up analyses revealed that the stimulus effect was driven mainly by an overall lower response to WN (WN < BOS, CON1, CON2, and REV-BOS), and a higher response evoked by CON1 (CON1 > REV-TUT; Tukey’s HSD, *p* < 0.02 for all stimulus comparison; all *post-hocs* were corrected for multiple comparisons here and throughout; ***Fig. 7B***). To avoid pseudo-replication (Picciotto, 2018), and because of the main effect of stimulus, we opted to perform follow-up analyses on just CON1 data for NCM. Follow-up analyses did not yield any significant differences between recording hemispheres for stimulus-evoked firing in FAD-treated (*F*_(1, 46)_ = 0.513, *p* = 0.478) nor aCSF-treated subjects (*F*_(1,47)_ = 0.734, *p* = 0.396).

**Fig. 7.**
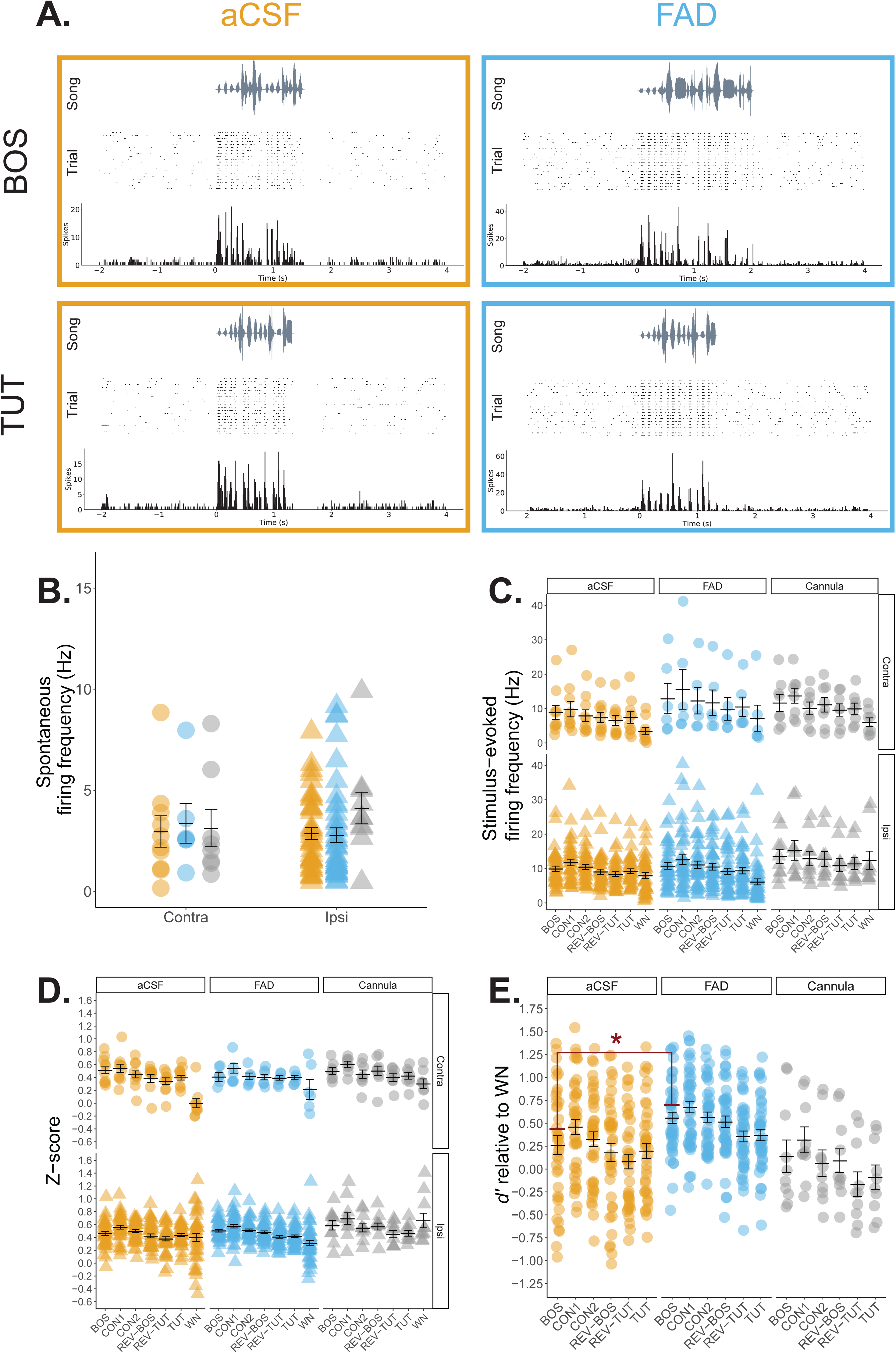
Single-unit recordings in NCM reveal modest differences in auditory responses in adulthood. **A**, Representative NCM single-unit recordings from an aCSF and FAD in response to presentations of birds’ own song (BOS) and tutor song. Each recording includes a song spectrogram (*Top*), and raster plot (*Middle*) with corresponding peri-stimulus time histogram in 10 ms bins (*Bottom*) across a 6 second period. The same unit is presented for each treatment across the two stimuli. **B,** Spontaneous firing rates were unaffected by developmental microdialysis treatment. Orange = aCSF; blue = FAD; grey = cannula; circle = contralateral hemisphere (relative to microdialysis site); triangle = ipsilateral hemisphere (relative to microdialysis site). **C**, Stimulus-evoked firing rates were significantly lower for WN and overall higher for CON1. A recording hemisphere x treatment interaction was significant; however, post hoc analyses limited to CON1 found no statistical differences for either treatment. **D**, Analysis of normalized auditory response (z-score) yielded a significant stimulus x recording hemisphere effect: contralateral NCM responded less to WN compared to all other stimuli, whereas forward conspecific stimuli elicited higher responses in the ipsilateral NCM, irrespective of treatment. **E,** Ipsilateral *d’* values relative to WN. BOS selectivity was higher in FAD songbirds in the ipsilateral hemisphere. BOS = birds’ own song; CON1; CON2 = conspecific song; REV-BOS = reverse bird’s own song; REV-TUT = reverse tutor song; TUT = tutor song. * = *p* < 0.05.

While raw firing rate data are informative, it is also useful to consider normalized auditory response rates (*z*-score) which accounts for recording site variability in spontaneous and stimulus-evoked activity (e.g. Vahaba et al., 2017). Analyses revealed a significant main effect of stimulus (*F*_(6, 651)_ = 17.643, *p* < 0.001) and recording hemisphere (*F*_(1, 651)_ = 12.935, *p* < 0.001), as well as a significant interaction between stimulus and recording hemisphere (*F*_(6, 651)_ = 3.051, *p* = 0.006; ***Fig. 7C***). In contralateral NCM, WN elicited a significantly lower z-score compared to all other stimuli (*p* < 0.001 for all stimulus comparisons). In contrast, z-scores were typically higher for non-reversed conspecific stimuli in the ipsilateral hemisphere regardless of treatment (CON1 > REV-BOS, REV-TUT, TUT, and WN; BOS > REV-TUT and WN; CON2 > REV-TUT and WN; *p* < 0.05 for all stimulus comparisons). Overall, the results in NCM suggest that irrespective of treatment, forward, conspecific stimuli (i.e. CON1, CON2, and BOS) reliably evoke the highest normalized auditory responses in the cannulated hemisphere.

Our initial impetus in recording from microdialyzed subjects was to test whether representations of learned stimuli (i.e. BOS and TUT) were different based on treatment early in development. To address this question, we calculated d prime (***d’***; see *Methods*) relative to WN to determine stimulus selectivity, as described in previous studies (Adret et al., 2012; Yanagihara and Yazaki-Sugiyama, 2016; Moseley et al., 2017). We limited our analyses to TUT and BOS as these were the learning-related auditory stimuli of interest that may have been impacted by treatment. Because of our earlier findings for auditory response profiles, we compared *d*’ scores separately by recording hemisphere. Treatment did not impact overall TUT selectivity for either contralateral (*F*_(1, 14)_ =2.222, *p* = 0.158) or ipsilateral (*F*_(1, 79)_ = 2.861, *p* = 0.095) recording sites in NCM. However, FAD subjects demonstrated significantly stronger BOS selectivity in the ipsilateral cannulated (*F*_(1, 79)_ = 6.371, *p* = 0.014; ***Fig. 7D***), but not contralateral hemisphere (*F*_(1,14)_ = 3.93, *p* = 0.067; ***Supp. Fig. 2A***). Taken together, unilateral E2 suppression in NCM during development enhances BOS representation in NCM relative to control birds.

#### HVC

The sensorimotor nucleus HVC contains a population of tutor-song-selective cells (Volman, 1993; Prather et al., 2008; Vallentin et al., 2016; Moseley et al., 2017) and receives E2-sensitive, indirect projections from NCM in part via the nucleus interfacialis of the nidopallium (Nif; Remage-Healey and Joshi, 2012; Pawlisch and Remage-Healey, 2015). To determine whether suppressing E2 synthesis in development affected downstream representations of either BOS or tutor song, we also recorded from HVC. Baseline firing rates were similar across treatments, recording hemisphere, and no interaction between the two factors were found (*F*_(1, 47)_, *p* > 0.132; ***Fig. 8A***). For stimulus-evoked firing, there was a main effect of stimulus (*F*_(6, 329)_ = 5.83, *p* < 0.001) and recording hemisphere (ipsilateral > contralateral; *F*_(1, 329)_ = 10.661, *p* = 0.001; ***Fig. 8B***). All other effects and interactions were non-significant (*p* > 0.131). Follow-up analyses revealed that, as expected, BOS elicited a significantly higher evoked firing response compared to all stimuli except TUT (BOS > CON1, CON2, REV-BOS, REV-TUT, and WN; *p* < 0.05); no other stimulus comparisons were significantly different.

**Fig. 8.**
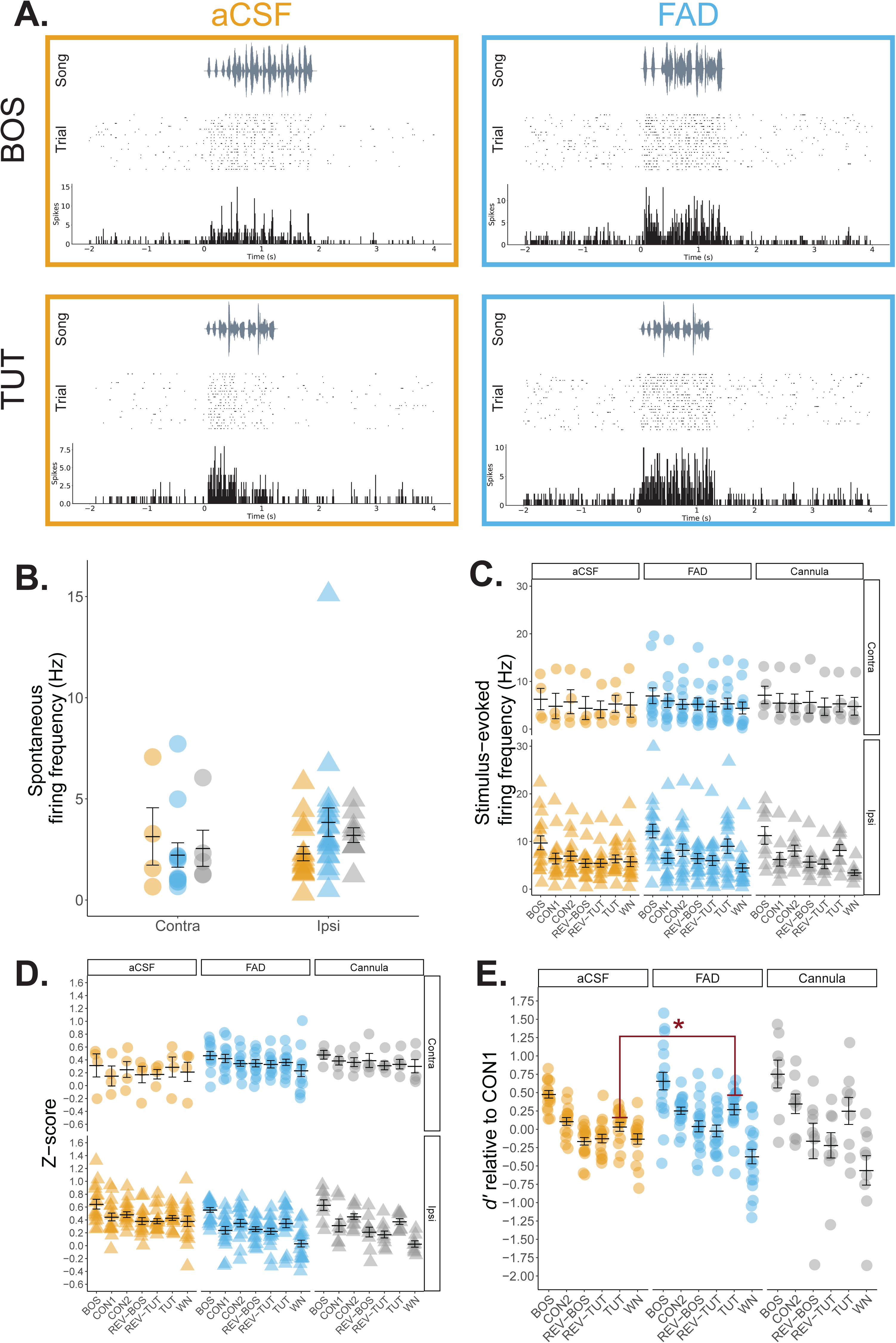
Tutor song selectivity is elevated in single HVC neurons of formerly estrogen-suppressed adult songbirds. **A**, Representative HVC single-unit recordings from an aCSF and FAD in response to presentations of birds’ own song (BOS) and tutor song. Each recording includes a song spectrogram (*Top*), and raster plot (*Middle*) with corresponding peri-stimulus time histogram in 10 ms bins (*Bottom*) across a 6 second period. The same unit is presented for each treatment across the two stimuli. **B,** Spontaneous firing rates were similar across treatments. Orange = aCSF; blue = FAD; grey = cannula; circle = contralateral hemisphere (relative to microdialysis site); triangle = ipsilateral hemisphere (relative to microdialysis site). **C**, Stimulus-evoked firing rates were significantly higher for BOS compared to all other stimuli except for TUT. Further, ipsilateral HVC displayed higher overall stimulus-evoked firing rates compared to contralateral HVC, independent of treatment. **D**, Analysis of normalized auditory response (z-score) yielded similar results as with firing rate; namely, a significantly higher response to BOS over all other stimuli independent of treatment, as well as a significantly suppressed response to WN compared to CON2 and TUT. **E,** Contralateral d’ values relative to CON1. TUT selectivity is significantly higher in FAD subjects solely in the contralateral hemisphere. BOS = bird’s own song; CON1; CON2 = conspecific song; REV-BOS = reverse bird’s own song; REV-TUT = reverse tutor song; TUT = tutor song. * = *p* < 0.05.

As with NCM, we also analyzed normalized auditory response in HVC. There was a significant effect of stimulus (*F*_(6, 329)_ = 10.384, *p* < 0.001), treatment (*F*_(1, 329)_ = 11.297, *p* < 0.001), as well as a significant interaction between recording hemisphere and treatment (*F*_(1, 329)_ = 25.745, *p* < 0.001; ***Fig.8***). All other main effects and interactions were non-significant (*p* > 0.176). As expected, BOS elicited a significantly higher response than did all other stimuli (BOS > CON1, CON2, REV-BOS, REV-TUT, TUT, and WN; *p* < 0.016). Conversely, HVC was less responsive to WN compared to select forward conspecific stimuli (WN < CON2 and TUT; *p* < 0.009). Based on the enhanced response to BOS for both z-score and stimulus-evoked firing, we opted to focus our follow-up tests on BOS. No significant differences were found for treatment for either the contralateral (*F*_(1, 14)_ = 1.097, *p* = 0.313) or the ipsilateral (*F*_(1, 33)_ = 1.223, *p* = 0.277) hemisphere.

For selectivity analyses, we focused solely on BOS and TUT relative to CON1 and tested whether TUT and BOS were differently represented between treatments. As there was a significant effect of stimulus and recording hemisphere, we analyzed the effect of treatment on TUT and BOS selectivity separately by hemisphere. BOS selectivity was statistically similar across treatments across both the ipsilateral (*F*_(1, 33)_ = 1.691, *p* = 0.202; ***Fig. 8D***), and contralateral hemisphere (*F*_(1, 14)_ = 0.804, *p* = 0.385; ***Supp. Fig. 2B***). In contrast, HVC units were more selective for TUT in the ipsilateral hemisphere of FAD subjects (*F*_(1, 33)_ = 5.82, *p* = 0.022; ***Fig. 8D***), but not contralateral hemisphere (*F*_(1, 14)_ = 3.45, *p* = 0.084; ***Supp. Fig. 2B***). Taken together, unilateral E2 synthesis inhibition in NCM enhanced the neural selectivity for tutor song in HVC, without causing changes in eventual tutor song imitation.

### Adult songbirds are unaffected by post-training inhibition of estrogen synthesis in NCM

As with juvenile songbirds, E2 is also acutely synthesized in the NCM of adult songbirds (Remage-Healey et al., 2008; Remage-Healey et al., 2012). Therefore, we also tested whether neuroestrogen production is involved in consolidating recent auditory experience in adult male zebra finches using a well-established auditory adaptation paradigm (see ***Methods***). Overall, adaptation rates (slope) were significantly shallower (lower) for familiar vs. novel stimuli (familiar = -0.28 ±0.4, novel = -0.49 ± 0.06; *F*_(1,4 122)_ = 4.150, *p* = 0.044), independent of treatment (*F*_(2, 122)_ = 1.182, *p* = 0.310) or an interaction between treatment and stimulus type (*F*_(2, 122)_ = 0.349, *p* = 0.706; ***Fig. 9***). Thus, unilateral estrogen synthesis in NCM immediately post-training did not adversely impact memory consolidation across development and in adulthood.

**Fig. 9.**
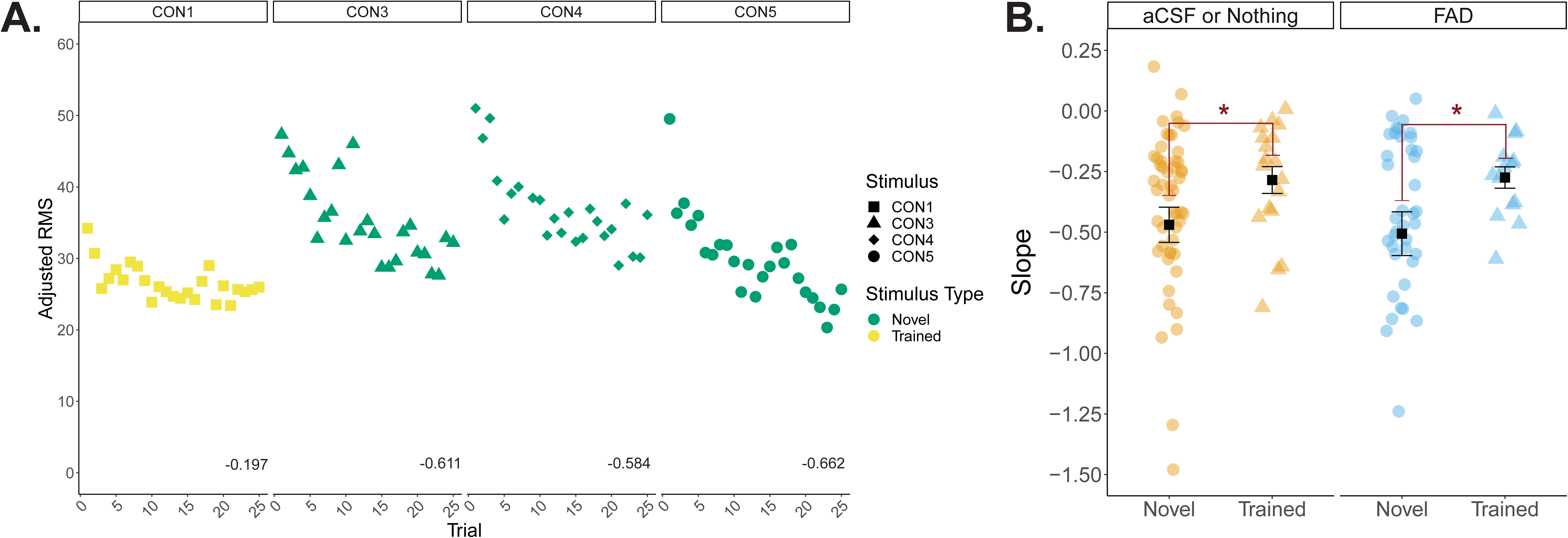
Neural adaptation to learned song is reduced in adult NCM independent of post-training E2 synthesis inhibition. **A**, An exemplar multiunit response in the NCM of an untreated hemisphere. Adjusted RMS declines at a faster rate (steeper slope) for novel song (CON3, CON4, and CON5) compared to a shallower slope (slower adaptation) for the recently exposed song (CON1). Slopes for each stimulus is shown at the bottom of each panel. **B,** Average slope per stimulus in aCSF or non-treated hemispheres compared to FAD-treated hemispheres in NCM; slope derived from multi-unit RMS. Orange = aCSF or no treatment; blue = FAD; circles = novel stimuli (three unique CON per bird); triangles = trained stimulus (a single unique CON). The y-axis has been compressed for clarity and five slope data points were omitted (-3.37, -2.72, -2.56, -1.58, 0.51). * = significant main effect of stimulus type (novel vs. trained); *p* < 0.05. CON = conspecific song.

## DISCUSSION

Our collective findings indicate that while aromatase is present in developing auditory cortex, global and unilateral attenuation of neuroestrogen production does not impair tutor song memorization. However, neuroestrogen blockade leads to suppressed singing rates during development and enhanced neural representations of tutor song in a downstream sensorimotor nucleus when measured in adulthood. Therefore, contrary to E2’s presence facilitating hippocampal-dependent adult spatial memory in a variety of species, downregulation of neuroestrogens may be permissive for auditory memorization, and its synthesis may be primarily important for exerting a pleiotropic effect on communication production and representation throughout the lifespan in songbirds. Taken together, this study is the first, to our knowledge, that tests the involvement of estrogen synthesis in consolidating an ethologically-relevant sensory memory within the developing auditory forebrain. Therefore, this study extends our knowledge of the role, region, and age in which estrogen is involved in learning.

### Developmental and regional shifts in neuronal cell density in NCM

We observed a decline in NCM cell density in sensorimotor-aged birds compared to sensory-aged subjects. Only one prior study, to our knowledge, has assessed the cell density of NCM across development and found no regional nor age differences in sensory-aged (20 and 30 dph) and adult male zebra finches (Stripling et al., 2001). It is unclear why our results diverge from those of Stripling et al. (2001), other than the resolution in the current study of sensorimotor vs sensory stages. Our findings suggest a form of experience-dependent network pruning that are consistent with heightened auditory responses in NCM in sensory- vs. sensorimotor-aged male songbirds (Vahaba et al., 2017). Alternatively, the volume of NCM many expand with age, leading to decreased neuronal density. To our knowledge, the volume of NCM across development has not been well characterized, and these ideas remain to be tested.

The density of cells in dorsal NCM was higher compared to ventral NCM, in contrast to previous observations (Stripling et al., 2001). This effect was independent of age, suggesting an anatomical distinction in developing NCM that may persist in adulthood (M. Macedo-Lima & L. Remage-Healey, unpublished observations). Numerous studies have described dorsal/ventral differences in response to auditory stimuli in NCM, but there does not appear to be a consensus regional effect. For example, immediate-early gene (IEG) auditory responses (i.e. *ZENK* induction in NCM in response to auditory playbacks) yield varying results depending on species: no differences between NCM subregions are reported in in adult male European starlings (Gentner et al., 2004) or adult male budgerigars (Eda-Fujiwara et al., 2012), whereas higher dNCM ZENK compared to vNCM has been reported in both adult female white-crowned sparrows (Sanford et al., 2010) and both sexes of adult black-capped chickadees (Phillmore et al., 2003; but see Avey et al., 2014). In contrast, extracellular recordings in the NCM of adult starlings find stronger experience-dependent changes in firing rates in ventral vs dorsal NCM (Thompson and Gentner, 2010), which were suggested to be attributed to a noted enhanced thalamic input from Field L to ventral NCM (Vates et al. 1996). Therefore, while subregion NCM divisions are anatomically distinct, the functional significance of this density difference across development is unclear, but are suggestive of regional differences in auditory responsiveness.

In addition to quantifying NCM neuronal density, we observed similar expression of aromatase and parvalbumin protein across the critical period for song learning. While aromatase expression has been assessed previously across development and in adults, we found that subregions within NCM of sensory- and sensorimotor-aged males possess a similar numerical capacity for estrogen synthesis. As aromatase is similarly expressed in NCM across development, changes in precursor androgens may explain previously observed age-dependent differences in baseline E2 in NCM across the critical period (Chao, et al., 2014), specifically in parallel with the maturation of the testes. Further, our findings with parvalbumin are in-line with recent findings that find that PV cell density is largely unchanged in across development in the NCM of in male and female zebra finches, as well as other auditory forebrain nuclei (Cornez et al., 2018). Therefore, PV-dependent inhibitory tone and estrogen production remain relatively unchanged across development, suggesting important roles throughout the juvenile period in males.

### Brain estrogen synthesis and song production in developing songbirds

Our experiments with systemic FAD treatment suggest that E2 facilitates song production in juvenile songbirds. It is well established that singing is regulated by classic (genomic) steroid hormone action, such as E2, in adult songbirds. In adult male zebra finches, long-term aromatase inhibition leads to suppressed courtship displays, including song production (Walters and Harding, 1988). More recent studies have found that neuroestrogen production also appears to acutely facilitate song production in adult zebra finches (Remage-Healey et al., 2009; Alward et al., 2016). Our data expand on this understanding that acute suppression of E2 production constrains singing to now include developing male songbirds. The neural locus of this effect of E2-withdrawal on song production is unknown, but likely to include social behavior network nuclei such as the aromatase-rich nucleus taenia (Saldanha et al., 2000; Ikebuchi et al., 2009). Androgens, namely testosterone, have classically been thought to be the critical hormone for the onset of motor production in developing songbirds. For example, plastic song emerges alongside the rise of testosterone in juvenile swamp sparrows (Marler et al., 1987). However, it has also been noted that circulating estrogens coincides with the onset of subsong (Marler et al., 1987). Thus, our data suggest that E2, and the conversion of precursor androgens to E2 within specific brain areas, may play a more significant role in song production in development than previously thought.

### Circulating and brain-derived estrogens are not required for tutor song memorization

Overall, systemic aromatase inhibition yielded minimal effects on eventual tutor song similarity. While tutor song similarity was slightly lower at 49 dph in FAD subjects, FAD-treated birds quickly ‘catch-up’ to comparable tutor song similarity levels as control birds, and produce songs of equal valence for adult female conspecifics. These results are novel given the relatively limited number of studies that have directly tested the role of hormones in song learning in male songbirds. Androgens are associated with the crystallization of plastic song (Korsia and Bottjer, 1991; Whaling et al., 1995; Bottjer and Johnson, 1997) and neural circuit development (Livingston and Mooney, 2001). In contrast, circulating estrogen levels are thought to promote plasticity due to their coincident rise in age-limited song learning in birds during the auditory memorization (“sensory”) phase of development (Pröve, 1983; Weichel et al., 1986; Marler et al., 1987; Marler et al., 1988; but see Adkins-Regan et al., 1990). While our sample size is limited, the data suggest that circulating estrogen synthesis is not required for tutor song memorization during development.

One important caveat for the systemic FAD experiment here is that our pharmacological treatment may have missed a putative ‘critical’ post-training consolidation period (e.g., within the first ∼30 mins following tutoring). E2 is important for auditory processing in adult and juvenile songbirds; thus, we did not want to interfere with online auditory processing of the tutor song during a tutoring session/playback. Instead, we intentionally administered FAD immediately after the offset of tutoring to specifically target the post-training memory consolidation period as in studies on hippocampal E2 and memorization (Frick, 2015). Comparable systemic aromatase inhibition treatments in birds led to marked reductions in E2 and aromatase activity (Wade and Arnold, 1994; Remage-Healey et al., 2010; Rensel et al., 2013; Alward et al., 2016), and systemic injections lead to suppressed aromatase activity in NCM within 30 mins (Alward et al., 2016; but see Krentzel et al., in submission). Thus, if systemic FAD actively suppresses E2 synthesis >30 minutes after administration, and the putative post-training auditory memory consolidation period is <30 minutes, it is important to consider that the pharmacokinetics of oral FAD may not sufficiently target the period of immediate post-training auditory memory consolidation.

In agreement with our systemic results, targeted unilateral suppression of neuroestrogen synthesis in NCM failed to prevent birds from eventually successfully imitating their tutor’s song. Tutoring leads to an initial drop in acute E2 levels within NCM, followed by a rapid increase immediately after a tutoring session in juvenile songbirds (Chao et al., 2015). In our paradigm, FAD was presented at the onset of tutoring and for a one-hour period immediately following the tutor session, without any detectable differences in eventual song similarity. Therefore, unilateral E2 synthesis in NCM does not appear to be required for auditory memory consolidation.

Additionally, juvenile songbird behavior is seemingly unaffected by unilateral estrogen manipulations in the auditory forebrain. Birds spent comparable amounts of time near by the live tutor and were similarly active during tutoring sessions. These results add to a small but growing understanding of tutor and pupil behavior during song learning. To our knowledge, these results are novel given the limited studies that explicitly quantify pupil behavior during tutoring (lab-reared, or otherwise) (Chen et al., 2016; Carouso-Peck and Goldstein, 2019). Juveniles are thought to preferentially learn from, and as an extension, imitate, more aggressive males who are mated or feed them early in development (Zann, 1996). While it is largely unknown how pupil behavior during tutoring affects song learning, one key behavior appears to be pupil ‘attention’ during tutoring (Chen et al., 2016). Our study found that unilateral E2 synthesis did not impact attention, as quantified by time spent near the tutor, thereby not interfering with song learning and imitation.

### Developmental neuroestrogen synthesis blockade has enduring effects on neural representation of autogenous and tutor song into adulthood

Suppressing E2 in NCM during development led to enhanced adult neural representations of the tutor’s song in HVC. HVC is a sensorimotor nucleus that dually represents both autogenous and tutor song in developing (Volman, 1993; Nick and Konishi, 2005b, a) and adult (Prather et al., 2010; Moseley et al., 2017) songbirds, and is necessary for song learning (Roberts et al., 2012). One possibility is that if neuroestrogen blockade reduces singing in microdialyzed birds as in our systemic experiments, there may be a ‘catch-up’ period that leads to enhanced salience, coding, or replay (Dave and Margoliash, 2000) of the social model’s song (tutor) once E2 synthesis inhibition is ‘released’ in NCM. Thus, our findings are consistent with other recent findings in swamp sparrows, in which HVC tutor- and BOS-selectivity is independent of vocal imitation accuracy in adulthood.

Interestingly, FAD treatments enhanced upstream BOS selectivity in NCM compared to control birds. Auditory forebrain neurons (including NCM) are typically selective for conspecific vocalizations over synthetic noises (e.g. tones) (Stripling et al., 1997; Stripling et al., 2001), and have been noted for having a subpopulation of BOS-selective cells (Janata and Margoliash, 1999; Grace et al., 2003; Amin et al., 2004; Yanagihara and Yazaki-Sugiyama, 2016). In particular, NCM contains experience-dependent tutor song and dual tutor song/BOS selective neurons during development (Yanagihara and Yazaki-Sugiyama, 2016). Auditory responses in NCM are rapidly modulated by estrogens in adult (Remage-Healey et al., 2010; Remage-Healey and Joshi, 2012) and developing zebra finches (Vahaba et al., 2017). Therefore, in agreement with our findings in HVC with tutor song, acute manipulations of E2 in NCM during development appear to be important for changing representations of birds’ own song as well.

It is worth noting that our treatments were presented unilaterally, and there is thus a strong likelihood that contralateral NCM can compensate for depressed E2 production in our study, leading to robust tutor song memory and proper song imitation in adulthood. While NCM appears to have lateralized function both natively (Moorman and Nicol, 2014), and with regards to E2 (Remage-Healey et al., 2010; De Groof et al., 2017), there is scant evidence for lateralized expression of aromatase (Saldanha et al., 2000; Ikeda et al., 2017). Relatedly, there is the additional possibility that either acute (microdialysis) or chronic (systemic) administrations may lead to homeostatic increases in aromatase production and/or activity (e.g. Saldanha et al., 2000), or upregulation of E2 from other sources (e.g. gonadal; adrenal). For example, estrogen-suppressed adult zebra finches have increased aromatase protein levels in the hippocampus, but not NCM (Saldanha et al., 2000). Lastly, it is possible that cannulation-induced injuries across control and FAD treated subjects obscured any potential differences in song learning outcomes. That is, since guide cannulae dissociated on their own, brain injury from the cannula may lead to similarly poor song learning outcomes as with FAD treatment. However, this possibility is unlikely to explain our findings as both microdialysis and systemically-treated birds yielded comparable song similarity rates in adulthood.

### Song learning is gated by experience

Our study also replicates and extends prior observations that experience with social partners, in addition to age, can regulate the closure of the critical period in songbirds. Importantly, the lack of song crystallization by 130 dph was independent of treatment, further emphasizing that unilateral estrogen synthesis in NCM does not participate in modulating critical period plasticity in contrast to androgens which prematurely crystallize song and related neural circuits (reviewed above). Others have also noted abnormal song in adulthood in lab-tutored songbirds (Eales, 1985, 1987; Morrison and Nottebohm, 1993; Slater et al., 1993; Jones et al., 1996; Zann, 1996; Deregnaucourt et al., 2013), and found similar changes such as dropped syllables, reduced syllable lengths, and increased stereotypy once abnormal singing birds were exposed to other adult males. Our work highlights the important limitation of controlled lab tutoring paradigms, namely that it is both quality and quantity of experience that dictate the closure of critical period song plasticity.

### Recent auditory experience consolidation is insensitive to estrogen synthesis blockade in adult NCM

Our results in adult animals build on a well-established paradigm in which recent auditory experience is encoded in adult and developing NCM (Chew et al., 1995; Stripling et al., 1997; Smulders and Jarvis, 2013; Miller-Sims and Bottjer, 2014; Ono et al., 2016). As in prior reports, we observed robust auditory recognition memory in the NCM of adults, regardless of treatment group. We find that auditory recognition in adult songbirds is unimpaired by unilateral inhibition of E2 synthesis post-training.

Repeated exposures of a single conspecific song leads to neural ‘recognition’ up 48 hours later (Chew et al., 1995), which is impaired when global estrogen production is dampened (Yoder et al., 2012). Our findings suggest that while E2 is important for spatial memory consolidation in the hippocampus of both birds and rodents (Frick, 2015; Bailey et al., 2017), as well as chemosensory memories in the olfactory bulb (Dillon et al., 2013), this role does not extend to auditory cortex. In rodents, E2 is rapidly upregulated in dorsal hippocampus immediately following an object recognition training session (Tuscher et al., 2016). In contrast, repeated song exposure in male and female zebra finches leads to immediate increases in estrogen levels which tapers off following song playback or social exposure cessation (Remage-Healey et al., 2008; Remage-Healey et al., 2012). Therefore, a lack E2 production following acoustic communication exposure in adults may explain the lack of a role for E2 in NCM for consolidating the auditory experience.

### Conclusion

Here, we demonstrate that estrogens exert a complex role in the auditory cortex of developing male songbirds. Our findings show the capacity to synthesize neuroestrogens remains high throughout development alongside substantial age- and subregion-dependent changes in NCM cell density.

Systemic estrogen synthesis blockade led initially to suppressed singing behavior in juveniles following tutoring. Further, while song memorization was unimpaired by acute inhibition of E2 production following training in developing and adult songbirds, early life E2 manipulations in auditory forebrain lead to altered neural selectivity of autogenous and tutor song in NCM and downstream HVC in adulthood, respectively. Taken together, this study expands our understanding of the role of brain-derived estrogens in learning and memory. Historically, studies on rapid E2 signaling and learning have been largely focused on adults and hippocampal-dependent learning. Therefore, in addition to continuing to study the role of brain-derived estrogen signaling across a diverse range of animals (Remage-Healey et al., 2017), it remains important to test its function across different ages (Gresack et al., 2007b, a) and brain regions.

## Supporting information

SuppFig1

SuppFig2

## ACKNOWLEDGEMENTS

From the Healey Lab, we thank Drs. Maaya Ikeda, Ben Pawlisch, and Catherine de Bournoville for help with microdialysis, Matheus Macedo-Lima assistance with electrophysiology analysis and immunocytochemistry and subsequent imaging, Olivia Li and Alex Rizzo for help with histology, and Christina Moschetto for quantifying female phonotaxis behavior. We also thank Dr. Colin Saldanha (American University) for the aromatase antibody, Dr. James Chambers for imaging assistance (UMass Amherst/Light Microscopy Core), and Drs. David Vicario and Mimi Phan (Rutgers University) for advice on analyzing habituation physiology data. This work was supported by the National Science Foundation IOS 1354906 (LRH) and a University of Massachusetts Graduate Student Dissertation Grant (DMV).

## SUPPLEMENTARY FIGURES

**Supp. Fig. 1 – Song changes in formerly microdialyzed subjects after exposure to adult male song at 130 dph. A,** Spectrogram examples from two aCSF and two FAD microdialysis subjects. *Top row*: song at 130dph; *Bottom row*: song at 6 weeks+ 130dph after subjects were exposed to conspecific adult males. **B,** Histogram demonstrating that treatment had no bearing on whether microdialysis subjects altered their adult song after exposure to conspecific males.

**Supp. Fig. 2 – Contralateral d’ selectivity in single NCM and HVC neurons. A,** NCM; statistical analyses were performed for TUT and BOS, only (see *Results*). Other stimuli plotted for visual comparison. Irrespective of stimulus, responses were significantly higher in aCSF treated subjects in both ipsilateral and contralateral NCM. **B,** HVC; statistical analyses were performed for TUT and BOS, only (see *Results*). Other stimuli plotted for visual comparison. Both TUT and BOS selectivity were statistically similar in contralateral HVC.

