## Supplementary figures and images for "Blocking neuroestrogen synthesis modifies neural representations of learned song without altering vocal imitation accuracy in developing songbirds"

### SuppFig1

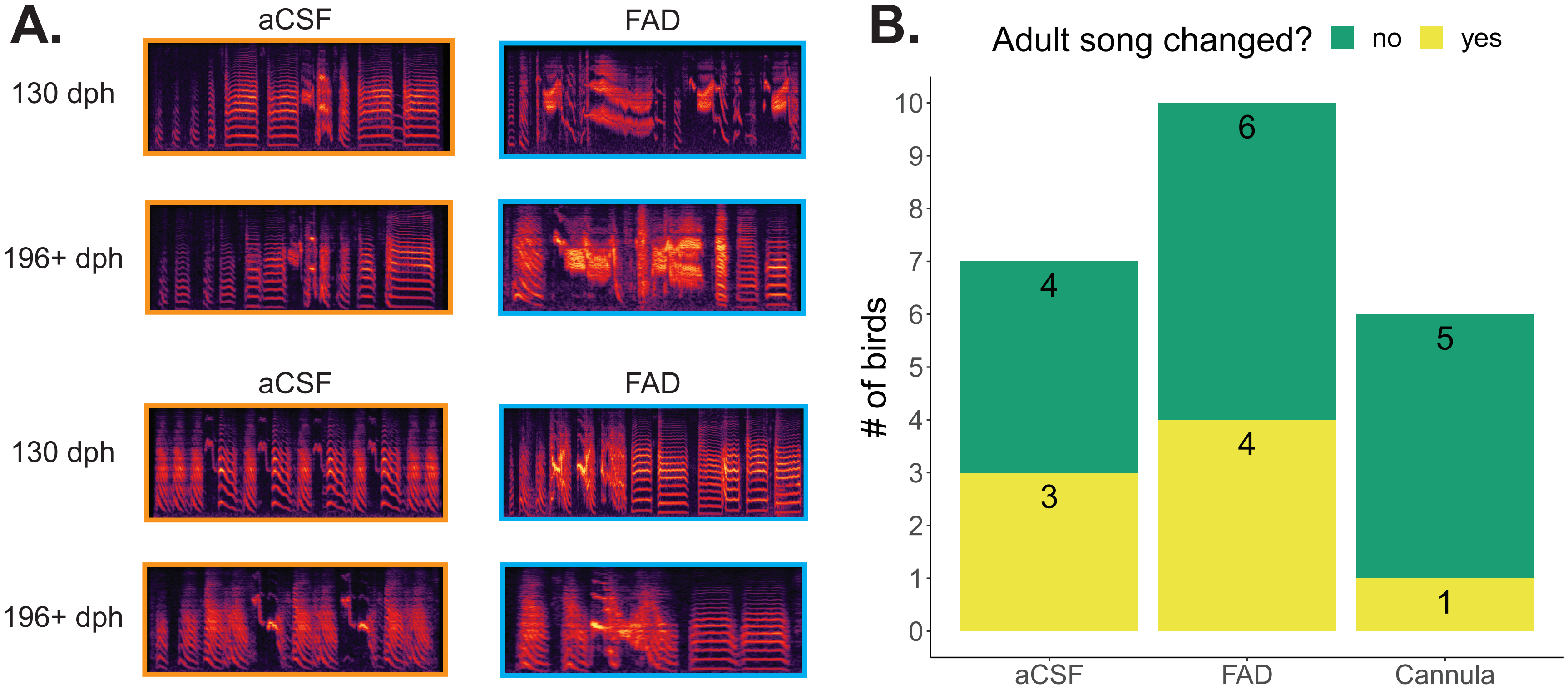

### SuppFig2

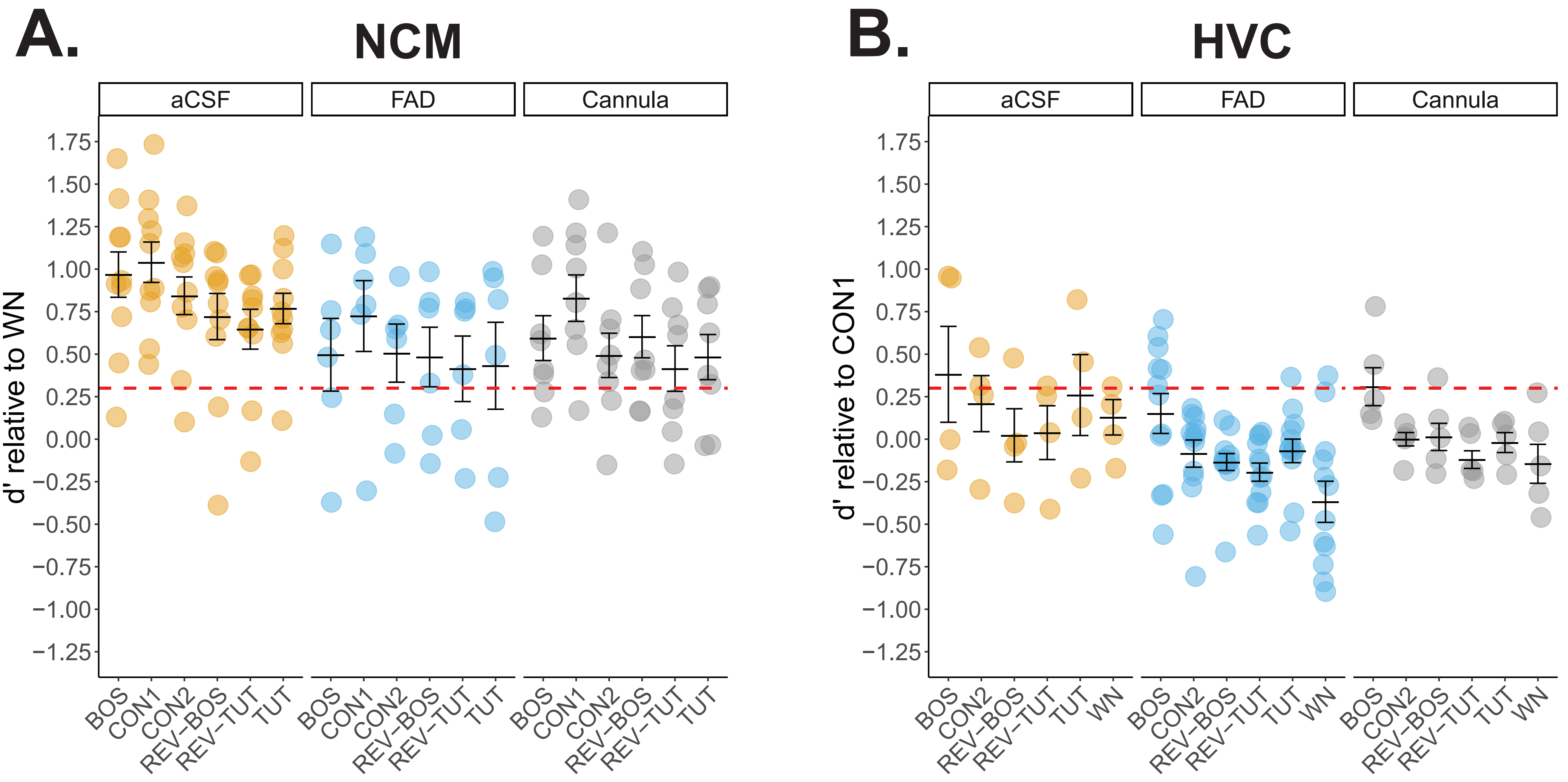
